# Misinterpreting electrophysiology in human cognitive neuroscience

**DOI:** 10.1101/2025.06.25.661032

**Authors:** Tzvetan Popov

## Abstract

An axiomatic view in contemporary neuroscience is that EEG components such as event-related brain potentials (ERPs) and oscillations are directly interpretable as manifestations of biological processes that support sensory, motor, and cognitive constructs of interest. This premise justifies and propels research programs in laboratories worldwide, but with a substantial social and economic cost, warranted by the potential for basic-science discovery and the resulting bench-to-bedside transfer for health and disease. But a different premise would be more fruitful. This article proposes that EEG components in psychophysiological experiments relate to cognition indirectly through their more direct relationship with oculomotor action.

The common experimental design that includes a baseline ocular fixation period preceding stimulus presentation provides an excellent template with which to develop the present proposal. Electrophysiological and eye-tracking evidence (3 published and 3 new data sets: 6 experiments, N_total_ = 204, in the context of face and affective picture viewing, reading, listening, rest, and microsleep) demonstrates how and why common conclusions, and reliance on them in clinical practice/treatment efficacy and drug development studies, are at best premature. Results indicate that the oculomotor system plays a mediating role between such EEG phenomena and cognition. Present evidence supports a complementary view of how EEG can shape the development of a broader thought horizon in psychophysiological theory and practice.

## Introduction

A century after Hans Berger’s seminal discovery, EEG is routinely used today to gauge the mind during work. Measurement of event-related brain potentials (ERPs) and oscillations are routinely used to encode, decode, and model psychology, and results are interpreted as the manifestation of brain mechanisms that support and implement psychological processes. This at times circular logic is enabled by the boundaries of the accepted set of theoretical and empirical premises that guide the interpretation of the observed results.

The present article makes a case for the role of oculomotor mechanisms as mediators in physiological measurement of cognitive processes. Such work falls squarely within cognitive neuroscience and thus within the broader field of psychophysiology (Marder & Miller, 2024). In psychophysiology (whether EEG, MEG, fMRI, pupillography, diverse cardiovascular measures, etc.), the assumption that under controlled experimental conditions the measured signal reflects the neural manifestation of mechanisms that allow direct inference of cognition justifies the use of EEG in psychology, with the expectation of uncovering the set of rules and relationships within and between brain circuits that enable the mind.

What does EEG measure? Dare we ask this question given the profound evidence of a century of work cementing the empirical observation that, irrespective of the sensory domain and experimental paradigm boundaries, the P1-N170 ERP sweep complex is generated upon stimulus presentation in the time domain, or followed by power modulation of oscillations in the time-frequency domains?

For the remainder of this narrative, I invite the interested reader to accept the premise that across species the central nervous system evolved to support the hosts’ overt action in the environment. Widely accepted, replicable experiments in the visual domain will serve as examples to evaluate the conjecture that ERPs and oscillations primarily reflect oculomotor action consequent upon which an indirect inference about the manifestation of cognition can be drawn. Yet ERP and oscillation data themselves do not provide direct evidence for this inference, because such data are mechanistically remote from cognitive processes. For reasons that will become apparent below, this conjecture is likely generalizable across sensory domains and relevant psychological constructs.

Most non-invasive event-related electro- and magnetoencephalographic visual experiments performed since Bergers’ seminal discovery have one feature in common. Independent of the research questions and psychological constructs under study, a monochrome background with a fixation target is utilized to record and model a baseline condition. A pervasive assumption is that such a baseline is neutral with respect to the stimulus and any experimental manipulation and hence qualifies as the background activity against which time and time-frequency data are expressed and subsequently modeled and interpreted. The tacit assumption is that maintaining fixation during baseline is a psychophysiological state (providing associated data) irrelevant to the interpretation of task-induced data. This assumption is premature, it leads to the misinterpretation, and its rejection will enable a complementary view. The problem with the assumption is that it foregrounds the existence of a truly resting (actionless) condition, during which the literal and metaphorical eye of the participant is resting, and does not provide any useful interpretation of the task data beyond serving as a baseline. Nothing is further from the truth.

The world has to move for us to see. And when the world is (quasi) stationary, the eyes have to move for us to see. Fortunately, the eye is in constant motion (Martinez-Conde et al., 2008; Martinez-Conde, Macknik, & Hubel, 2004; Peter H. Schiller & Edward J. Tehovnik, 2001). That oculomotor action creates a redundancy of the sampled object, what would be eventually classified by the observer as a particular feature, such as edge, orientation, etc. The mere radiation of light from an object towards the retina when both the world (e.g., bar stimulus) and the retina are static does not result in a neural response. It is the relative motion that elicits a measurable pattern. In the seminal discovery by Hubel and Wiesel in anesthetized and paralyzed cats (hence no eye movements), single-cell firing was not recorded before “Suddenly, just as we inserted one of our glass slides into the ophthalmoscope, the cell seemed to come to life and began to fire impulses like a machine gun. It took a while to discover that the firing had nothing to do with the small opaque spot— the cell was responding to the fine moving shadow cast by the edge of the glass slide as we inserted it into the slot” (p. 61 in (Hubel & Wiesel, 2004)). An experimental approach utilizing “hand-held search stimuli” (p. 7734 in (Movshon & Newsome, 1996)) or a reversal beam ophthalmoscope that involves movements due to the nature of the reversal beam ophthalmoscope (Van Essen, Newsome, & Maunsell, 1984) uses a moving beam of light to illuminate different parts of the retina. It is the movement that allows for dynamic examination of the retina, enhancing the ability to observe details and identify abnormalities, given full anesthesia and a paralyzed participant. This strategy paved the way to record visual cells sensitive to edges, motion, and color properties of a stimulus in the anesthetized and paralyzed primate by moving and/or changing the occurrence of the stimulus over the targeted retinal position and the corresponding receptive field (e.g. (Essen & Zeki, 1978; Majaj, Carandini, & Movshon, 2007; Maunsell & Van Essen, 1987; Movshon & Newsome, 1996; Van Essen et al., 1984)). Contemporary technology might replace hand-held search movement and/or ophthalmoscope, yet the general approach that requires relative action to infer vision remains. Why and how is this action relevant for the interpretation of data acquired in non-invasive human psychophysiology?

In such research, the maintenance of fixation is considered a rest condition in which the eyes do not move, and therefore eye-muscle control is not a relevant or covarying feature that would require further examination and or interpretation of the acquired EEG data. Instead, it qualifies as a baseline condition. However, this assumption is wrong. Instead, as the default state of the eye is movement, maintaining fixation requires active control - not a true resting state. As will be outlined below, this active control contributes to the manifestation of stimulus-induced ERPs and oscillatory power modulations. A consideration of the circuits involving eye movement control will be helpful in understanding how these arise.

During the awake daytime, the eye is aligned along the so-called optical axis, which is 23° nasally from the orbital axis (Figure 1), the latter being the principal direction of eyeball orientation in the absence of eye muscle activity. Thus, the position that would qualify as true rest is 23° away from central fixation. From empirical, first-person experience, the position of the eye during the awake state is straight ahead 0° on the optical axis, always ready to evaluate the environment. However, this position is not resting. Miniature eye muscle contractions maintain a fixation position by continuous movements around a given target (Figure 1B) (Martinez-Conde et al., 2008; Martinez-Conde et al., 2004). These fixational eye movements are instantiated and controlled by the same circuit controlling other types of ocular action such as vergence, pursuit, and saccadic eye movements. The circuit provides eye muscle control enabled by neurons in the brain stem nuclei, superior colliculus (SC), and cortex. Except for the brain stem, the coding operation of deep-layer neurons in SC and cortex foregrounds an intricate relationship. Effects of stimulation of deep pyramidal neurons by a microelectrode in SC, visual, parietal, or frontal cortex converge on the observation that independent of where the current eye position is, the same vector saccade is produced, with identical vector direction and amplitude (Figure 1) (P. H. Schiller & E. J. Tehovnik, 2001). Neither the rate nor the frequency of the microelectrode stimulation determines the initiation of the saccade. Instead, a window of approximately ∼100-120 ms acts as a “reader mechanism” within the temporal boundary of which a single saccade with the same size and direction is produced (Figure 1). Two saccades within ∼200 ms, 3 within ∼300 ms, and so forth (Robinson, 1972; Schiller & Stryker, 1972). This discovery has been termed a vector code or vector coding operation of the neurons and is distributed across the cortex as well as the SC (P. H. Schiller & E. J. Tehovnik, 2001). It aligns with the fact that the fastest movement in the human body is the saccade reaction time, which is experimentally confirmed to be no faster than ∼100 ms (Kingstone & Klein, 1993; Schiller, Sandell, & Maunsell, 1987). The maximum bursting rate of SC, frontal eye fields (FEF), and parietal cortex neurons occurs at ∼100 ms after stimulus presentation and aligns with the reaction time of the corresponding saccade (Stine, Trautmann, Jeurissen, & Shadlen, 2023; Zhu, Zhou, Constantinidis, Salinas, & Stanford, 2024). Cooling of deep-layer pyramidal neurons in the primary visual cortex (V1) eliminates saccades towards a stimulus, and temperature restoration restores the saccadic behavior identical to that before cooling (Schiller, Stryker, Cynader, & Berman, 1974). Similarly, cooling the FEF introduces a failure to maintain fixation on a target, while cooling SC is associated with an increase in saccade reaction time and reduction in saccadic amplitude (Keating & Gooley, 1988). Ablation of just two structures, SC and FEF, eliminates all visually guided, that is stimulus-induced, eye movements (Schiller & Tehovnik, 2005) and truncates the participant’s overall ocular range (Keating & Gooley, 1988). This type of eye movement control, i.e. maintaining fixation and directing eye position toward a target, is an overarching requirement built into experimental designs using fixation maintenance as a baseline in human non-invasive psychophysiology. Invasive electrophysiology in non-human primates has shown that these eye movements follow a ∼100 ms temporal code. How and why is this experimental observation relevant to the interpretation of data in non-invasive human psychophysiology?

**Figure 1:**
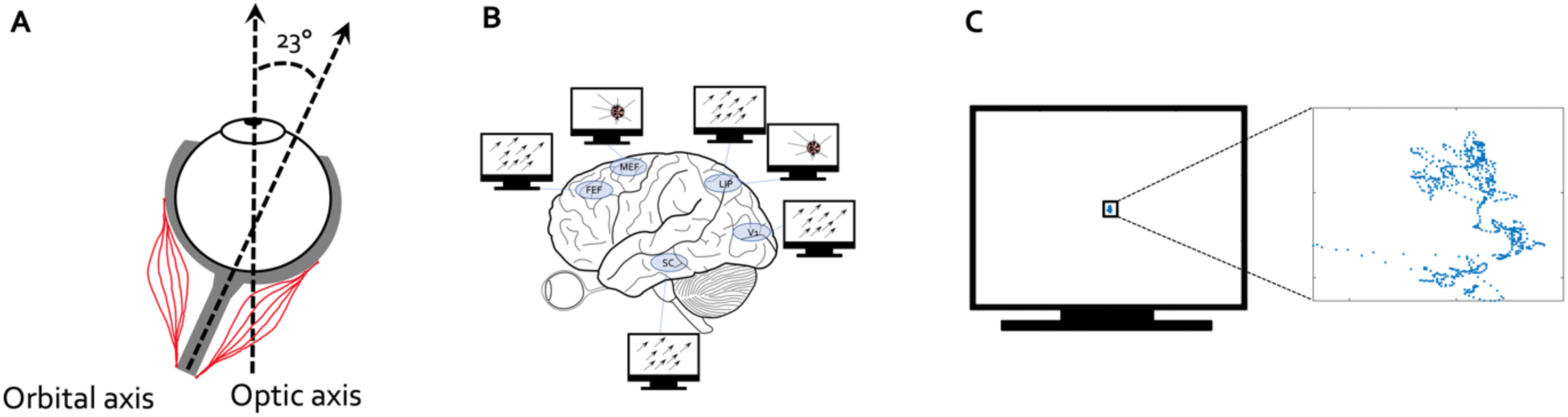
The default state of the eye is movement. **A-** Ocular muscles support the continued maintenance of eye positionalong the optic axis (during the awake state and in the absence of other eye movements such as saccades, vergence, and pursuit). **B-** Experiments in non-human primates utilizing microelectrode stimulation of pyramidal cells in deep cortical and SC layers confirm a coding operation directing the eye along a vector of a fixed size and direction independent of current eye position. This vector code depends on a temporal window of ∼100 ms independent of duration or frequency of the electric stimulation. Neurons in the parietal cortex (e.g., lateral intraparietal area LIP in the non-human primate) and the medial eye field (MEF in the non-human primate, or the premotor cortex in humans) exhibit also motor-fields: independent of the current gaze position, the eye is brought into a particular position on the visual display upon electric stimulation of the corresponding neurons. C-Maintenance of fixation during cognitive experiments entails miniature, yet preserved ocular action. It is an action typically not noticed ing EOG and below a predefined eye movement (e.g., micro- or macro-saccade) threshold but equally engaging and requiring cortical circuit control involved in the manifestation of A and B.

In the absence of a microelectrode “pacemaker”, an oscillation with a duty cycle of ∼100 ms could, in principle, substitute for the microelectrode stimulation and the temporal window discovery mentioned above. The collective dynamic of pyramidal neurons residing in the cortical deep layers (e.g., the local field potential LFP) has been associated with oscillations in the alpha 8-14 Hz frequency range. That is, it is conceivable that an ongoing duty cycle brings enough neurons together such that their net firing aligns with a given principle direction and directs the saccade toward an object, much like the microelectrode stimulation does. Such cooperation is a requirement because the ongoing maintenance of the brain’s internal dynamic interferes with single-neuron input-output integration and subsequent firing, which is artificially overcome during microelectrode stimulation.

Such intricate neuronal relationships are worth pondering but help little in the interpretation of non-invasively recorded event-related potentials and fields. At best, they provide grounds for discussion and interpretation that could easily lead to speculation. And yet, the ∼100 ms width of a P1-N170 sweep complex or ERP is often casually interpreted by researchers as the “neural underpinning” of perception, attention, distractor suppression, and social cognition (e.g., face processing) without eyebrows being raised.

The N170 ERP component was recently characterized as the “first psychiatric biomarker” in the FDA’s Biomarker Qualification Program selected for use in the “enrichment of clinical drug development trials” in children diagnosed with autism spectrum disorder. That is, an ERP component has been judged to be an objective measure of social functioning [for a recent discussion see e.g. (Floris et al., 2025; Mason et al., 2022)]. The latter is arguably one of the more complex cognitive constructs we study. But social cognition cannot happen that quickly after face stimulus presentation. Similarly, the presentation of an emotionally arousing visual scene “indicate[s] selective processing of emotional stimuli, reflecting activation of motivational systems in the brain” (Cuthbert, Schupp, Bradley, Birbaumer, & Lang, 2000). This is a strong statement presented as a solid fact, publicized in textbooks (Gross, 2007) and clinical trial practice as an indicator of and basis for inferences about cognitive-behavioral treatment efficacy (Garland, Froeliger, & Howard, 2015; Greimel et al., 2020; Zsigo et al., 2024). ERPs are interpreted readily (and in effect directly) as reflecting “cognitive” phenomena (e.g., face and/or affective processing) and should instead be interpreted directly as an oculomotor control action, with cognitive consequences as important epiphenomena.

The fact of the matter is, though, that the ∼100 ms full-width half maximum of the P1-N170 sweep complex is closer to the observations concerning oculomotor action. First, neuronal populations in the cortex utilize a 100 ms temporal window to initiate an eye movement; second, these neurons produce a collective rhythm (i.e., alpha oscillation) with a 100 ms duty cycle which in turn affects their output and therefore eye movement; and, third, the fastest saccadic reaction time is ∼100 ms.

Above, I outlined some established features of the operation of the neural systems that control ocular position and motion, and I suggested that some prominent interpretations of ERP components as direct readouts of cognitive processes are unwarranted, because a better interpretation is that those ERP phenomena directly reflect neural control systems rather than cognition. The present contention is not that cognition is unrelated to such ERP components or that cognition should not be inferred from them but that a critical role of 0cular control systems is left out of such thinking. In fact, the presence of the P1-N170 sweep complex informs the experimenter about the termination of fixation during baseline and the direction of the eye toward aspects of the stimulus. The former is not "cognition” itself but the necessary ocular action required to visually explore the latter. This active visual exploration is reflected in the presence of later ERP components and in power modulation of alpha oscillations. Consequently, action or overt behavior can be understood as having the purpose of controlling perception (Powers, 1973), as an overarching principle about what the organism is trying to accomplish. Everything is behavior. What is meant by ”behavior” in the present narrative is the type of behavior that motivates and justifies psychophysological research such as perception, attention, and (working) memory. This type of behavior is controlled by oculomotor action.

Below, I provide a series of empirical observations derived in the context of widely accepted and utilized psychophysiological experiments such as passive viewing of face stimuli, complex visual scenes, listening to speech, and reading to support the conclusion that ERP components and oscillations often primarily reflect oculomotor action and the neural control systems that support it. Thus, the present contention is that some interpretations of EEG phenomena do not attend sufficiently to relevant neural control systems driving behavior, do not understand that oculomotor action in large part to manage perceptual input, and that these phenomena occur as part of a cybernetic control system. Thus, such interpretations do not accurately attribute the direct role of some EEG phenomena to the correct node or level in the control systems.

## Materials and Methods

### Participants

A total of 167 participants were recruited across the three human group studies reported here: the face perception experiment, the picture viewing experiment, and the word reading experiment. These studies were not preregistered. The sample size was limited by resource constraints, such as time constraints related to the duration of the courses within which the experiments were conducted, as well as budget limitations. As argued by Lakens(Lakens, 2022), the potential limitations are acknowledged here, given the practical constraints preventing the achievement of an ideal sample size based on power analysis. Participants were recruited from the local university and through community advertisements. The sample comprised 126 female and 41 male participants, with an age range of 18 to 72 years (M = 26.3 years). In the face perception experiment, 55 participants took part (43 female; age range: 20–45 years; M = 25.9 years). In the picture viewing experiment, 84 participated (65 female; age range: 18–44 years; M = 24.7 years). In the word reading experiment, 28 volunteered (18 female; age range: 18–72 years; M = 31.9 years). Prior to participation, all participants provided written informed consent in accordance with the Declaration of Helsinki. The studies were approved by the local ethics committee at the host institution. Participants received either monetary compensation or course credit for their participation.

Additionally, publicly available data were used. Details regarding the procedures for the story listening experiment (featured in Figure 14) can be found in (Armeni, Güçlü, van Gerven, & Schoffelen, 2022), information about the resting-state data and procedures is available in (T. Popov, Trondle, et al., 2023), and details on the microsleep data are available in in (Hertig-Godeschalk et al., 2019; Skorucak et al., 2019).

### Acquisition Setup and Procedures

In all experiments, visual scenes, faces, or words were displayed on a SyncMaster P2770HD monitor with 1920×1080 pixel resolution. Participants were seated in front of the monitor at an approximate distance of 70 cm, positioned in a chin rest. Stimuli were presented using PsychoPy (Peirce et al., 2019) on a full-screen grey background.

EEG data were acquired using the mBrainTrain Smarting device, a wireless 24-channel EEG system. The system utilizes semi-dry/saline-based Ag/AgCl electrodes embedded in a cap, adhering to the International 10–20 System of electrode placement. Electrodes were referenced online to the FCz electrode, with AFz serving as the ground. Impedance for each electrode was maintained below 10 kΩ. Data were acquired at 250Hz sampling rate and converted offline to an average reference for further analysis.

Eye-tracking data were acquired using a Tobii Pro Fusion eye tracker, with real-time gaze data transmitted via the Lab Streaming Layer (LSL) protocol. The eye tracker captured left and right gaze points and pupil diameter, streaming data at a 250 Hz sampling rate. Triggers marking stimulus onset were sent via LSL using a separate stream for synchronization of the EEG and eye tracking streams.

### Task Design

The face task was adopted from previous reports (e.g.(T. Popov, Miller, Rockstroh, & Weisz, 2013; T. G. Popov, Rockstroh, Popova, Carolus, & Miller, 2014)). Briefly, prior to the beginning of the experiment, a fixation cross appeared on the screen for 5 s before the first trial presentation. Each trial consisted of a 5 s video presentation of a face morphing from neutral to a fearful or happy expression of the same actor, or to another neutral expression of a different actor. Participants viewed a randomized sequence of the videos, each belonging to one of three emotion-based conditions, with 120 trials (40 per condition) presented. Each trial followed the sequence of a baseline fixation cross displayed for 3 seconds (±0.5 s jitter), followed by the video presentation for 5 s. The experiment continued until all videos had been presented. Valence and arousal aspects of the design have been evaluated elsewhere (T. Popov et al., 2013; T. G. Popov et al., 2014; Popova et al., 2014) and are not further analyzed here. The face images were derived from the Radboud Faces Database (RaFD, www.rafd.nl) including the stimulus examples illustrated here (e.g. Figures 2,3).

**Figure 2:**
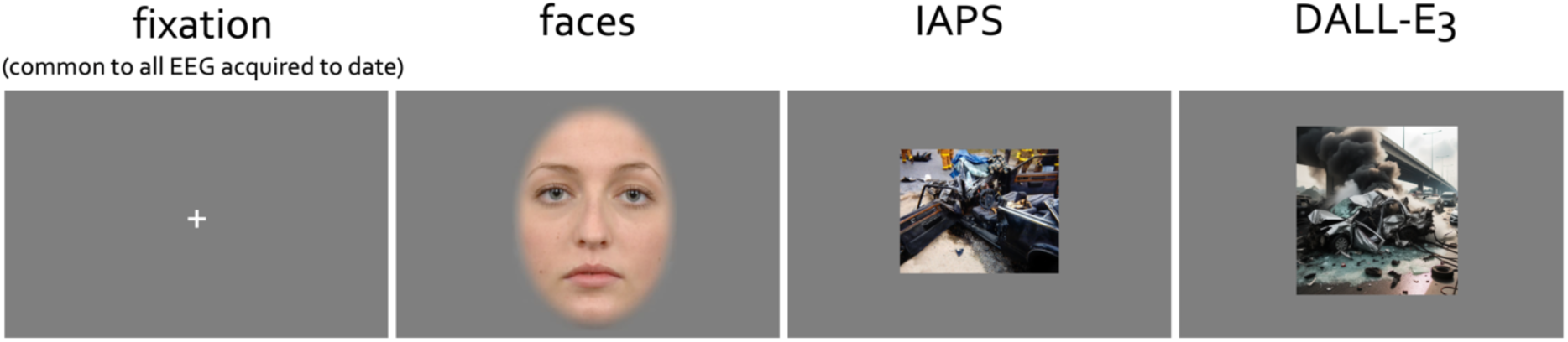
Experiments in human non-invasive psychophysiology often start with visual fixation on a minimal stimulus. It is used as a baseline and a contrast to all subsequent analyses. In the present narrative examples utilizing human face stimuli, emotionally arousing and neutral images from the IAPS database as well as images generated by artificial intelligence (DALL-E3) are reported. The duration of fixation viewing across experiments was at minimum 1 s, viewing duration of faces 5 s and of images 1 s. The face image is a stimulus example from the Radboud Faces Database (RaFD, www.rafd.nl).

**Figure 3:**
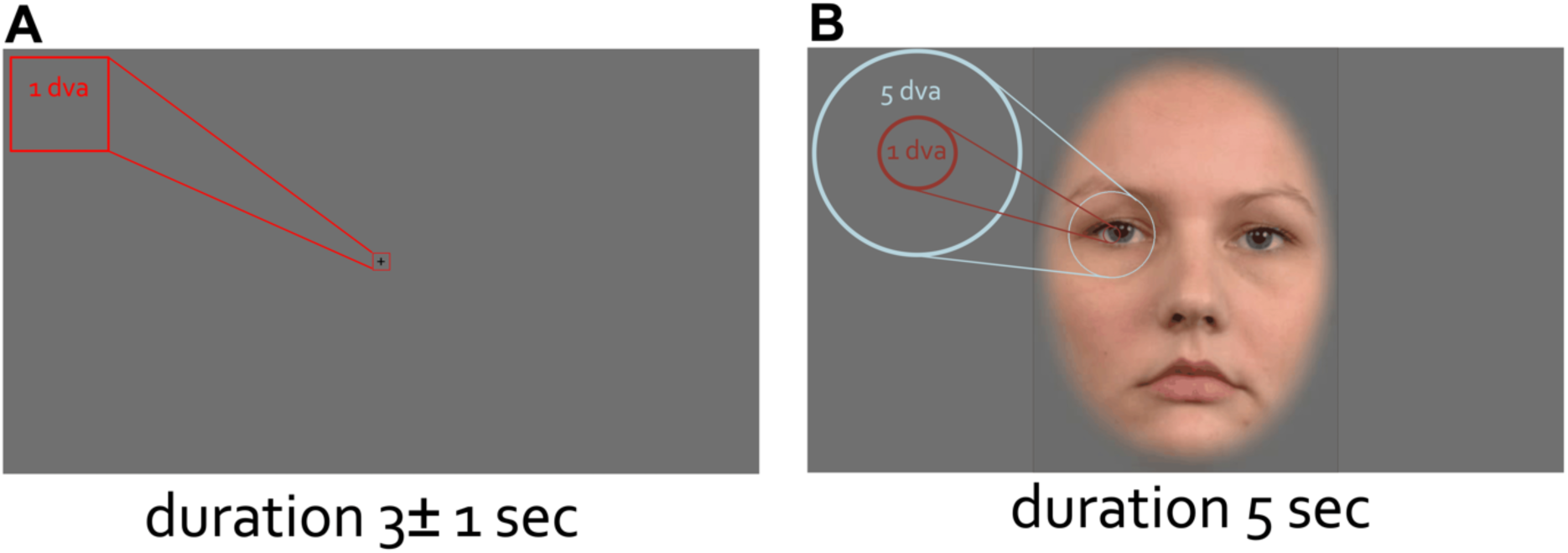
The size of the stimuli at scale. Fixation cross (A) and the face stimuli (B) at the original scale. Participants were positioned at 70 cm distance from the computer screen. The duration of the fixation viewing was at minimum 2 s and face viewing 5 s. The face image is a stimulus example from the Radboud Faces Database (RaFD, www.rafd.nl).

The picture viewing experiment consisted of multiple trials, with each trial structured as follows. Each trial began with the presentation of a fixation cross (+) on a black background for 5 s before the first stimulus appeared. Participants viewed images categorized into three experimental conditions: 100 pleasant, 100 unpleasant, and 100 neutral. Within the pleasant and unpleasant conditions, half of the images were sourced from the International Affective Picture System (IAPS) database, and the other half were pleasant and unpleasant images generated by OpenAI’s DALL-E 3. All 100 neutral images were from the IAPS database. This manipulation allowed for the examination of LPP effects elicited by images that neither have been systematically rated and evaluated with respect to valence and arousal nor are clearly identifiable as artificially generated. Each image was displayed at the center of the screen for 1 s, followed by a 1 s baseline with a ±0.5 sec jitter. Images were randomly shuffled for each session to ensure counterbalancing. In the pleasant and unpleasant condition 50 images were randomly chosen for each subject. Image IDs from the IAPS data base were as follows. **Pleasant**: 1710, 1720, 1722, 1811, 2045, 2058, 2075, 2155, 2160, 2209, 2216, 2345, 2346, 4006, 4007, 4090, 4180, 4220, 4225, 4250, 4255, 4290, 4310, 4320, 4490, 4505, 4520, 4525, 4530, 4533, 4542, 4597, 4599, 4601, 4606, 4607, 4608, 4609, 4610, 4611, 4612, 4619, 4623, 4625, 4626, 4641, 4645, 4651, 4652, 4653, 4656, 4658, 4659, 4664, 4666, 4668, 4670, 4680, 4687, 4690, 4693, 4694, 4695, 4698, 4800, 5470, 5600, 5621, 5623, 5626, 5628, 5629, 5660, 5700, 5825, 7330, 7405, 7499, 8001, 8021, 8030, 8031, 8040, 8041, 8158, 8179, 8180, 8185, 8186, 8190, 8200, 8210, 8260, 8370, 8400, 8420, 8490, 8492, 8496, 8500. **Unpleasant**: 1033, 1050, 1051, 1052, 1120, 1200, 1201, 1205, 1300, 1301, 1525, 1930, 1932, 2095, 2345.1, 2661, 2730, 2811, 3001, 3010, 3030, 3051, 3059, 3061, 3071, 3080, 3100, 3101, 3102, 3103, 3110, 3120, 3130, 3140, 3180, 3181, 3185, 3195, 3212, 3213, 3215, 3220, 3225, 3266, 3301, 3350, 3530, 3550, 6212, 6213, 6231, 6250, 6260, 6312, 6313, 6315, 6350, 6510, 6520, 6540, 6550, 6560, 6563, 6570, 6571, 8230, 9040, 9042, 9043, 9075, 9160, 9185, 9253, 9301, 9325, 9340, 9410, 9412, 9413, 9414, 9421, 9423, 9430, 9495, 9570, 9571, 9590, 9592, 9594, 9596, 9599, 9622, 9635.1, 9900, 9901, 9902, 9904, 9910, 9920, 9921. **Neutral**: 2002, 2102, 2191, 2200, 2210, 2214, 2273, 2305, 2308, 2357, 2377, 2381, 2383, 2390, 2393, 2396, 2397, 2411, 2480, 2484, 2493, 2495, 2499, 2512, 2516, 2570, 2595, 2749, 2840, 2870, 2880, 2890, 4233, 5471, 5500, 5520, 5530, 5533, 5534, 5731, 5740, 6150, 7000, 7002, 7003, 7004, 7006, 7009, 7010, 7011, 7012, 7014, 7016, 7017, 7019, 7025, 7026, 7032, 7034, 7035, 7038, 7041, 7043, 7045, 7052, 7053, 7055, 7056, 7057, 7059, 7061, 7062, 7080, 7090, 7100, 7130, 7150, 7175, 7179, 7185, 7186, 7187, 7217, 7233, 7235, 7255, 7287, 7491, 7493, 7512, 7513, 7547, 7550, 7595, 7705, 7710, 7950, 8312, 9210, 9700.

### Data Processing and Analysis

Offline data analyses were performed using the FieldTrip (Oostenveld, Fries, Maris, & Schoffelen, 2011) open-source toolbox for neuroelectric data analysis. Continuous EEG and eye-tracking data were segmented around the events of interest (stimulus onsets) into epochs, including a 2 s baseline prior to each trial’s stimulus onset.

A finite impulse response (FIR) band-pass filter (1-45 Hz) was applied to the raw data before artifact rejection. Independent component analysis (ICA) was then used to remove oculomotor and cardiac artifacts. Identified components corresponding to these artifacts were removed, and the cleaned ICA components were projected back into the raw, unfiltered data. After ICA correction, a low-pass filter (45 Hz) was applied to the reconstructed EEG data, ensuring minimal distortion by 50 Hz mains artifact while preserving low-frequency neural signals.

EEG data were further inspected for bad channels, which were interpolated using the distance option: electrode locations were aligned to the MNI standard brain using a head model, and neighbouring electrodes within 100 mm were identified for interpolation. If any channels were missing from the dataset due to hardware or signal quality issues, they were interpolated based on available neighbouring channels using FieldTrip’s ft_channelrepair function.

For time-frequency analysis, power estimates were computed using multitaper convolution with a Hanning taper and a fixed window length of 0.5 s, resulting in a 2 Hz frequency resolution for the range 2–40 Hz. The window was stepped every 50 ms across the epoch, covering the range of -2 to 5 s (in the face experiment) and -1 to 1 s (in visual scenes and reading experiments) around stimulus onset.

Power estimates were averaged across trials for each experimental condition separately. A baseline correction was applied using the -Inf to -0.25 second pre-stimulus interval, transforming the power values into decibel (dB) change relative to baseline.

The 1/f aperiodic component was removed from each trial using the *specparam* algorithm, which provide parametrization and visualization of periodic components in the continuous data distinguished from aperiodic activity (Donoghue et al., 2020). Similar time-frequency analyses were applied to the story listening and resting state/microsleep data, with adjustments made to the epoch length to match the specific temporal dynamics of each condition. The same multitaper convolution method was used, ensuring consistency across analyses.

### Gaze Data Processing

The raw eye-tracking data were analyzed by computing 2D gaze density heat maps and eye velocity estimates, providing a comprehensive characterization of visual scanning behavior.

For each time point in the respective dataset, gaze positions along the horizontal (x) and vertical (y) axes were selected. To compute gaze density, the data was binned into a 1000 × 1000 pixel grid, and a 2D histogram was created using MATLAB’s *histcounts2* function. The raw binned gaze data were then smoothed using a Gaussian filter (*imgaussfilt* with a smoothing factor of 5) to generate a continuous heat map representing gaze density over time. Following 2D transfrmation, the gaze heat maps were converted into a FieldTrip-compatible structure with: smoothed gaze density values stored in *powspctrm*, akin to spectral power in time-frequency analyses, and horizontal positions (time) and vertical positions (freq) as axes in the coordinate grid, ensuring compatibility with FieldTrip’s statistical and visualization functions. This transformation allows gaze density data to be analyzed using the same statistical approaches as time-frequency EEG/MEG data, facilitating direct comparisons between gaze behavior and neural activity.

### Eye Velocity Estimation

To examine eye movement dynamics, instantaneous gaze velocity was computed based on the horizontal and vertical eye position signals. The raw data were extracted and smoothed using a 100 ms moving average window (movmean over 25 samples). Velocity was estimated by computing the temporal derivative of smoothed gaze position, scaled by the sampling rate. The resulting absolute velocity values were stored in a FieldTrip-compatible structure. These preprocessing steps facilitated the integration of gaze position, gaze density, and eye movement velocity into a unified pipeline for statistical comparisons between eye-tracking metrics and neural oscillations across experimental conditions.

### Statistical Analysis

Statistical significance was assessed using the cluster-based permutation framework to control for multiple comparisons (Maris & Oostenveld, 2007). This method accounts for dependencies across electrodes, time points, and (where applicable) frequency by identifying spatiotemporal clusters of significant effects. A two-tailed alpha threshold of 0.05 was applied, and statistical significance was determined using 500 permutations.

### Data and Code Availability

All data and the code necessary to reproduce the present results are available at [https://osf.io/q4mez/].

## Results

In the following, complementary considerations supported by empirical observations are outlined. The mix of results and discussion sections, although atypical for research papers, is intentional to help the reader contextualize how past considerations motivate a re-evaluation and a complementary view of the results presented.

Reiterating the premise, the default state of the eye is movement, and the purpose of action, including eye movement, is to control perception. During awake states, the eyeball in orbit is aligned along the optical axis, maintained in this position by continuous and ongoing eye muscle activity (Figure 1A) (Wright, 2006). Eye movement is supported by vector coding operations of cortical and subcortical (SC) neurons (Figure 1B) (P. H. Schiller & E. J. Tehovnik, 2001), and persists during periods of fixation (Figure 1C) (Martinez-Conde et al., 2004).

Under this premise, ERP components and oscillations are examined first in the context of passive viewing of faces experiment (Figure 2). After a period of fixation lasting 2-3 s, a dynamic face stimulus, gradually changing expression, is presented for 5 s (details reported in (T. Popov et al., 2013)) but not necessary for the present discussion. Relevant here is that participants were tasked with maintaining fixation for some duration before evaluating a face stimulus. Participants sat at a distance of approximately 70 cm from the visual display. Given the size of the face stimuli, the fixation cross spanned approximately 1 degree of visual angle (dva) (Figure 3A), and the eyes and mouth sections spanned 5 dva (Figure 3B). These dva values were chosen given that compromised foveal vision results in difficulties in interpreting changes in facial expression (Lerner et al., 2006; Seiple, Rosen, & Garcia, 2013). From the observer’s perspective, compromised foveal vision makes it difficult to notice, infer, and label changes in facial expressions. For instance, if only part of the nose falls within foveal vision at 170 ms post face onset, interpreting an aberrant N170 as a marker of social cognition impairment in autism is questionable. The key features needed to infer socially relevant cues, such as the eyes, eyebrows, and mouth, have not yet been explored.

The arrangement of the fixation cross relative to the size of the visual display and the participant’s distance was consistent across all experiments reported here.

First, a single participant viewed the presentation of faces (120 trials) over 19 recording sessions, resulting in a total of N_trials = 2023 after artifact correction. Figure 4 illustrates the time course of the ERP averaged across occipital electrodes O1 and O2, along with the time course of eye velocity extracted from continuously recorded eye-tracking data. Additionally, the gaze position aggregated across 500 ms intervals is depicted at the top of the corresponding time series.

**Figure 4:**
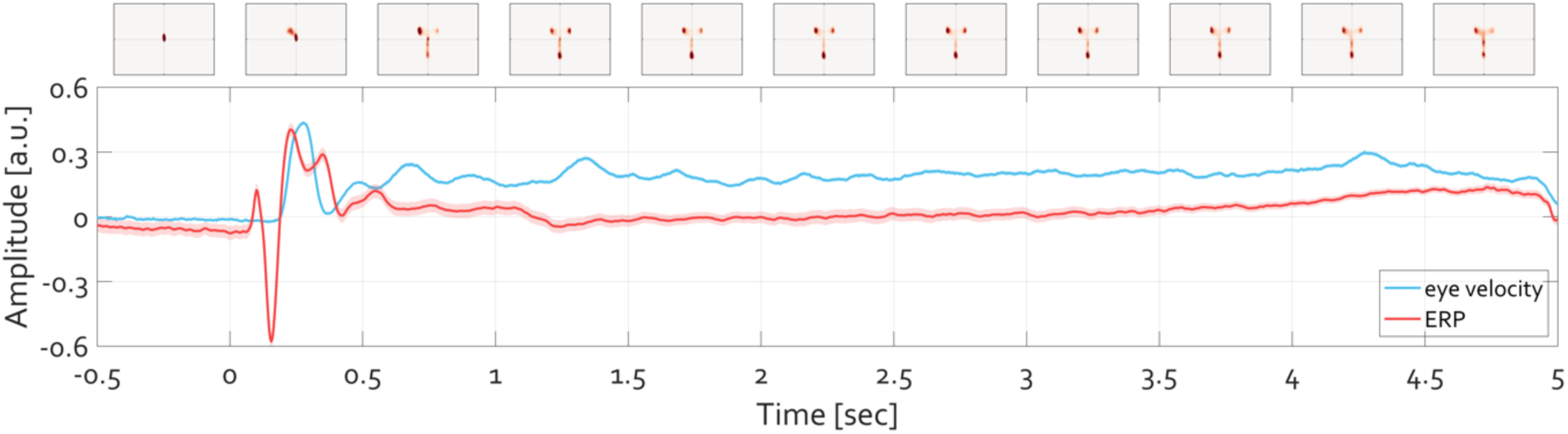
Time course of ERP (red) and eye velocity (blue). After initial fixation interval of 2 s (last 0.5 s shown) a face stimulus was presented for 5 s. The ERP averaged over occipital electrodes O1 and O2 is plotted in red. The shading around that tracing shows the standard error of mean (SEM) across trials (N_trials_ = 2023). The velocity of the eye deviating from the experimentally defined fixation location is illustrated in blue, with the shading corresponding to its SEM. The units (arbitrary unit a.u.) of ERP amplitude and eye velocity are range-corrected in order to facilitate comparison. The position of gaze per 500 ms time windows is illustrated at top, with heatmaps indicate the histogram of eye gaze position-the stronger the red color, the more frequent the location of gaze at the respective position.

Descriptively, during the baseline fixation interval, the eye remains positioned at the fixation location defined by the experimenter, with minimal deviation as indicated by the variance of the eye velocity (Figure 4 blue line). Following face presentation, a transient ERP (Figure 4, red line) and an increase in eye movement speed are apparent, continuing throughout the trial. The heatmaps at the top of Figure 4 summarize the 2D histogram of gaze locations visited by the eye averaged over 500 ms intervals. Most gaze locations are centered on the fixation during baseline, whereas during the trial gaze lands predominantly on the eyes and mouth in the face stimulus (Figure 4, top panels).

One can zoom in on the first 600 ms after the stimulus presentation to closely evaluate the time series (Figure 5). This examination reveals two insights. First, the change in eyeball rotation within the orbit occurs only after the P1-N170 sweep complex. In fact, the negativity aligns with the latency of N170 (Feuerriegel, Churches, Hofmann, & Keage, 2015). Second, the heatmaps quantifying the 2D histogram of gaze positions during the latency period of 0 to 170 ms are nearly identical to those during the baseline period (-170 to 0 ms). Thust, the shift in gaze position away from fixation (blue color in the heatmaps in Figure 5) toward face features (red color in heatmaps in Figure 5) becomes apparent only after the N170 component. Given the boundaries of foveal and parafoveal vision (Figure 3), at the time of what is commonly interpreted as the "face-selective" N170, it is implausible that the eyes have already encoded social-cognition-relevant features such as variations in eye and mouth parts.

**Figure 5:**
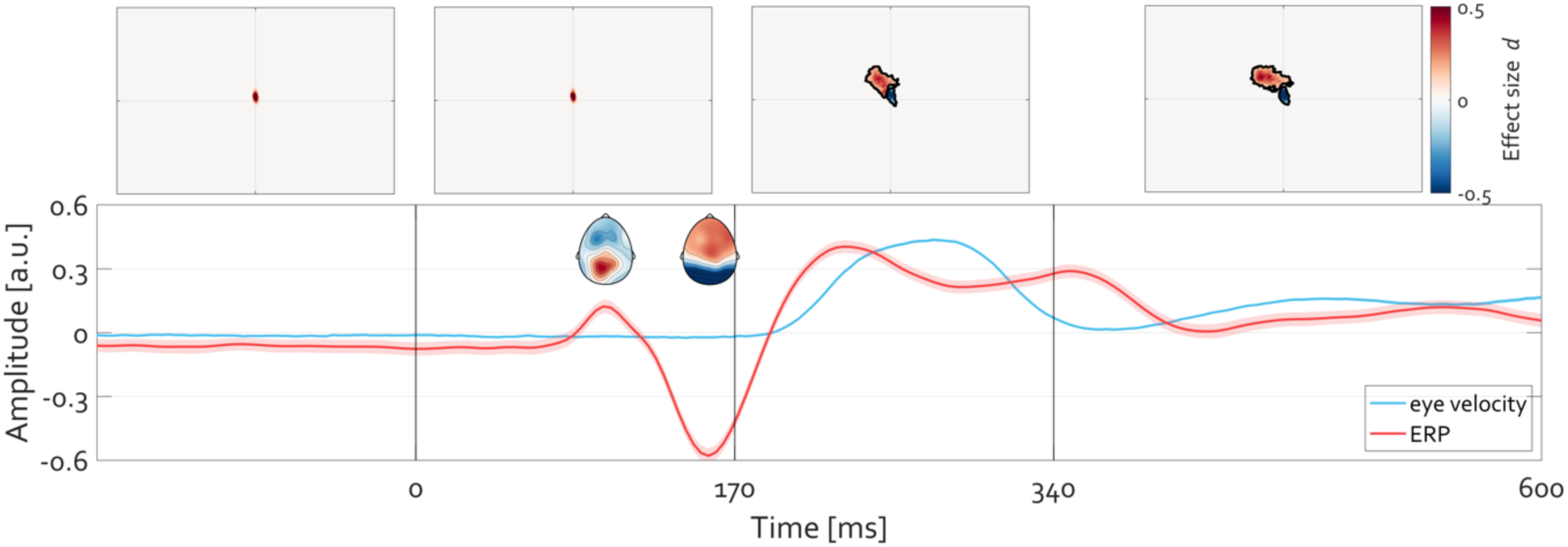
Initiation of eye movement toward face-specific features of the image follows the P1-N170ERP sweep complex. The time courses are identical to those reported in Figure 5. The inset topographies illustrate the corresponding topography of P1 and N170, respectively. A statistically significant (cluster permutation test corrected for multiple comparisons, p<0.05) deviation in gaze position is observed after the N170, indicating termination of fixation toward, in this case, the left eye of the actor in the stimulus (red color in the heatmap denoting positive Cohen’s d effect size). The aggregated position of gaze for the time interval 0 to 170 ms is nearly identical to that extracted from the baseline (-170 to 0 ms).

The EEG scalp topography around the P1 and N170 latency is the basis for inference about the underlying cortical generators and is interpreted as the electrophysiological basis of mechanisms implementing psychological constructs such as early visual perception and feature encoding. However, based on the findings just reviewed, this interpretation must derive from factors (e.g., mental pattern completion) other than a mere reflection of light radiating in straight lines from low-level features on the screen, passing the lens and radiating further within the vitreous humor within the eyeball without site damage due to the complexity of light diffusion in liquid, passing the layers of retinal neuropil without further site damage, and eventually reaching and innervating the photoreceptor cells. I.e., N170 cannot be a downstream manifestation of face perception as is commonly assumed. At a minimum the transition of features in the displayed face stimulus would require that these features be actively visited by the observer’s eye. They were not.

This conclusion was replicated at the group level. N_group_ = 55 individuals performed the same task (Figures 6 and 7). Latencies of the early transient ERP components were very similar to those observed in the single individual. Again, a transient increase in eye velocity followed N170 (Figure 6). During the fixation period, the heatmaps depicting the 2D histogram of gaze locations were centered around the center of the visual display and then tracked the contours of a face throughout the trial (Figure 6, top panels). Zooming in on the first 600 ms replicated that, before the P1-N170 sweep complex, eye movements did not visit the facial features of the actor. Locations corresponding to eye and mouth features were visited by the participant’s eyes only evident after the N170 (Figure 7). The topographic difference of the N170 component in the single individual case (Figure 5) vs. the group (Figure 7) is not material to this conclusion, as the difference is likely due to variability of in position of the standard 10-20 electrode placement relative to variability in the underlying gyrification and anatomy (in the group data) vs. the relatively low variability in the single participant case.

**Figure 6:**
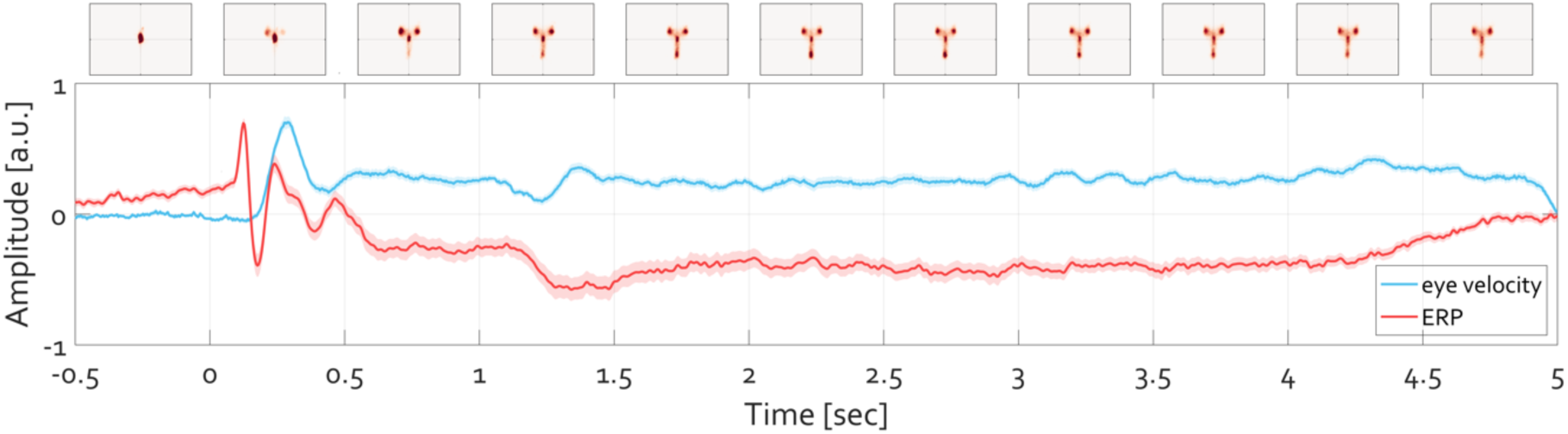
Grand average (N_group_=55) of the time course of ERP and eye velocity. The line colors and design of the illustration are identical to those of Figure 4.

**Figure 7:**
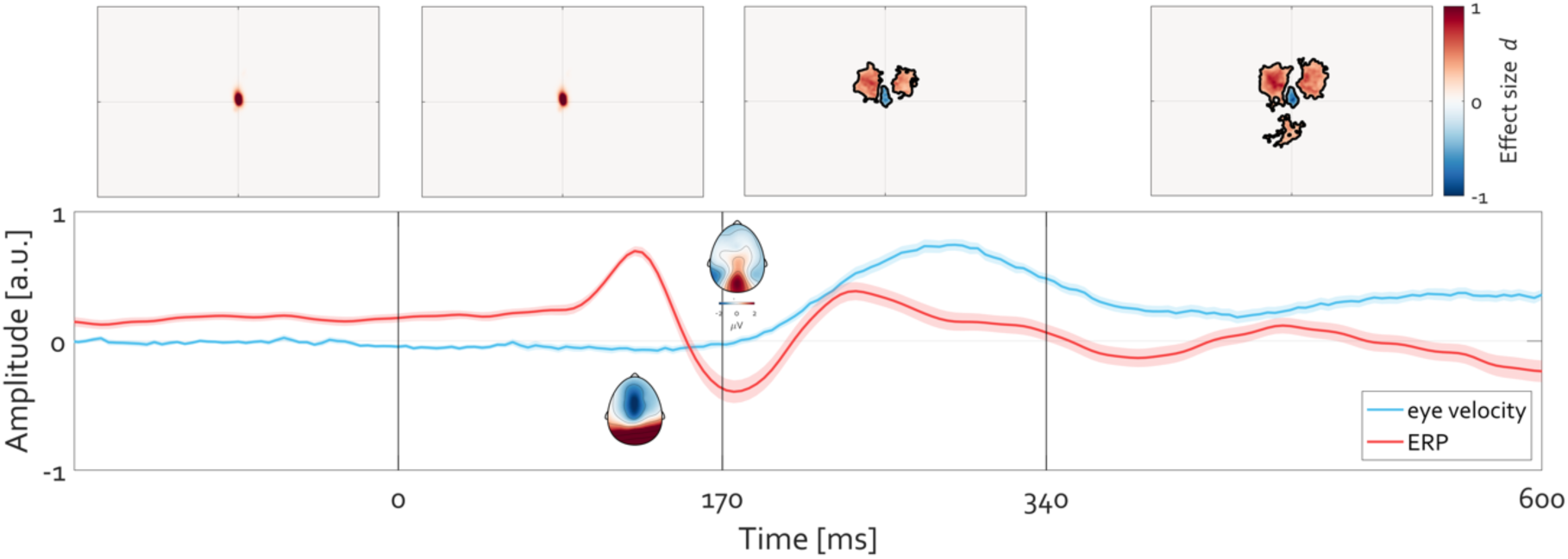
Grand-average illustration of the initiation of eye movements. **On group level (N_group_ =55), eye movements** toward face-specific features of the image follow after the P1-N170ERP sweep complex. Line colors and design of the illustration are identical to Figure 5 but illustrate a group result rather than a single participant.

Beyond ERP waveforms, the oculomotor account of human alpha oscillations (Tzvetan Popov, Miller, Rockstroh, Jensen, & Langer, 2021) emphasizes a relationship between the power modulation of alpha oscillations and the direction of gaze, as well as eye exploration more broadly. Empirical evidence supports the conclusion that higher exploration through eye movements correlates with a stronger decrease in alpha amplitude and vice versa (T. Popov, Gips, Weisz, & Jensen, 2023; T. Popov & Staudigl, 2023).

To assess the generalizability of this relationship in the context of face evaluation, the 2D histogram of gaze positions was computed aggregating over consecutive 500 ms intervals, and a slice approximately ± 5° vertical dva around the central fixation was extracted. A sliding window (500 ms) was used to visualize the time-resolved maintenance and departure from fixation (Figure 8). When juxtaposed with the time-frequency-resolved activity averaged over occipital electrodes (O1, O2), the relationship between increased alpha power and fixation maintenance becomes evident (Figure 8). Descriptively, maintenance of eye position (against the default state) on a fixation target coincides with higher occipital alpha amplitude than with eye exploration during face viewing. As will become clear shortly, leaving this at a descriptive value for now brings a benefit by considering stimuli lacking face features first.

**Figure 8:**
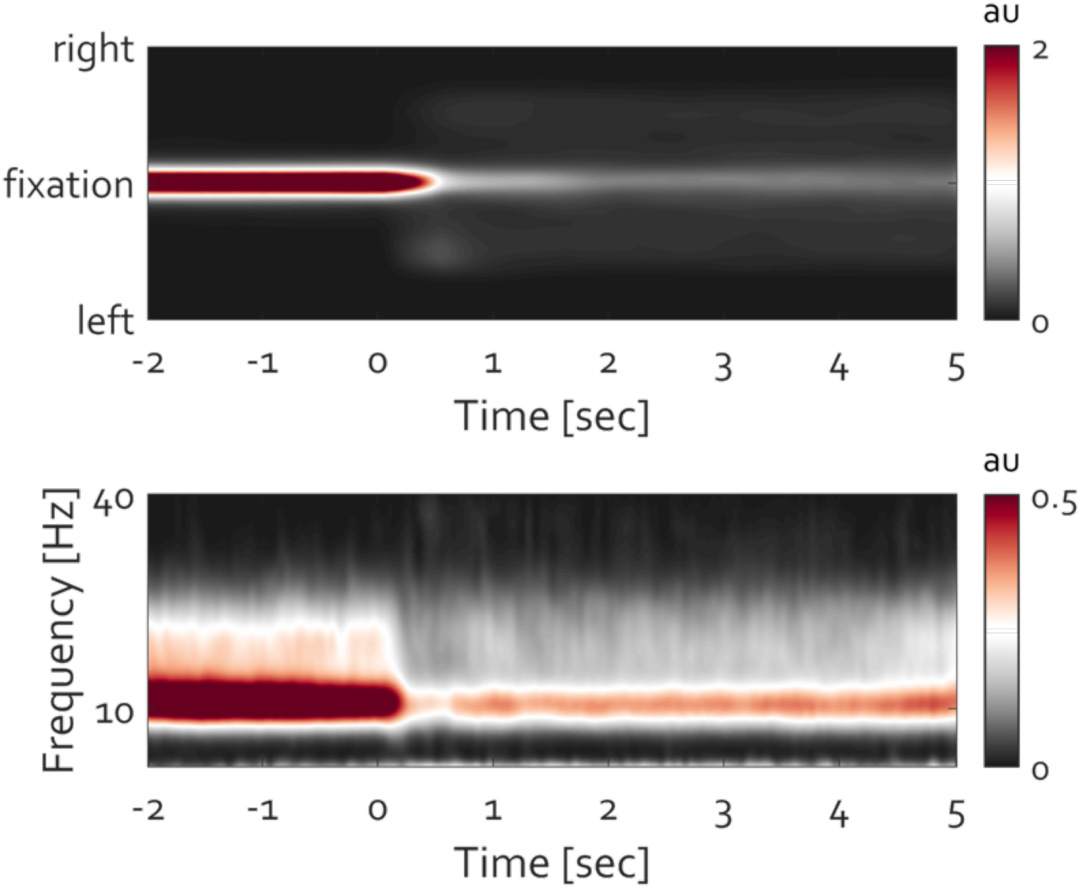
Occipital alpha power modulation and maintenance of fixation are related. **Top- time-resolved** histogram of gaze density around fixation (ordinate). The red color highlights increased density of gaze positioning at fixation. Bottom-the corresponding time course of frequency-resolved activity over electrode ‘POz’. Warm colors indicate the strength of oscillatory power (expressed in arbitrary units). The time course of power modulation corresponds to the time course of fixation maintenance. Departure from fixation toward exploring the face stimulus is associated with reduction of alpha power in line with previous reports (Popov et al. 2022, Popov and Staudigl 2023)

The relationship between eye exploration and evoked brain potentials and oscillations was evaluated in the context of viewing complex scenes. Images from the IAPS database, including 100 neutral, 50 pleasant, and 50 unpleasant images with high arousal ratings, as well as 50 pleasant and 50 unpleasant images generated by artificial intelligence using DALL- E3, were utilized (image IDs and ratings of IAPS images as well as the AI images used are accessible via https://osf.io/q4mez/). Participants viewed a total of 300 images presented pseudo-randomly for 1 second each. The baseline period was jittered between 1-2 seconds, during which participants were instructed to maintain central fixation.

The single participant evaluated in Figure 5 also viewed these images (300 in total from the IAPS database only) across 24 sessions, resulting in N__trials_ = 5496 after artifact correction. Figure 9A illustrates the ERP averaged across parieto-occipital electrodes (’P3’, ’Pz’, ’P4’, ’POz’). Following the initial P1-N170 sweep complex, a positive slow wave known as the late positive potential (LPP) was observed, which was stronger for arousing images compared to neutral ones (pleasant and unpleasant conditions combined). The topographies depict the distribution of activity during baseline, at the N170 latency, and the difference in topography (arousing minus neutral) during the LPP latency from 400 to 1000 ms post-stimulus (often defined and examined as a difference wave hence the subtraction) (Cuthbert et al., 2000). This evaluation confirms the well-documented hedonic arousal effects in the ERP literature (Hajcak & Foti, 2020; Schupp et al., 2000).

**Figure 9:**
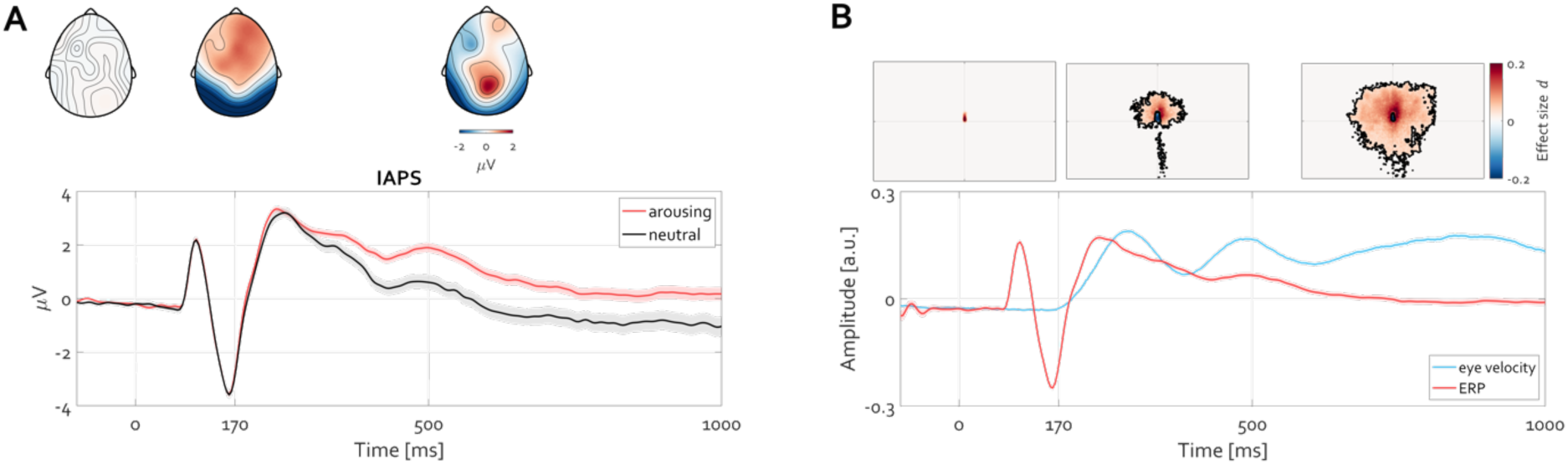
Initiation of eye movement toward features of a visual scene follows after the P1-N170 ERP sweep complex. **A-** ERP illustrating the initial P1-N170 sweep complex followed by a slow potential commonly referred to as the late positive potential (LPP-stronger for arousing compared to neutral images). The inset topographies illustrate the corresponding topography during the baseline, N170, and the difference (arousing minus neutral) for the LPP latency 400-1000 ms post-stimulus onset. **B**-ERP time course pooled across both conditions (red line) and the corresponding eye velocity (blue line). Shading denotes SEM across trials. The heatmaps at top illustrate the 2D histogram of gaze positions from stimulus onset until 170 ms after (left), the statistical contrast baseline (-170 to 0 ms) vs. 170-500 ms, and the statistical contrast baseline vs. 500 to 1000 ms post visual scene onset. Statistical testing was carried out with cluster permutation tests corrected for multiple comparisons (p<0.05) expressed in units of Cohen’s d effect size.

Examining the time course of eye velocity (Figure 9B), a relationship nearly identical to that observed during face viewing was present. The temporal progression of the P1-N170 sweep complex was followed by an increase in eye velocity. Heatmaps confirmed that the single individual examined above terminated fixation instructions and began exploring details in the visual scene, yet again only after the N170 latency, very similar to viewing faces (e.g., Figure 5).

This relationship was confirmed at the group level (N__group_ = 85). Figure 10A illustrates grand-average activity across all participants and parieto-occipital electrodes. The grand-average time course of eye velocity evolved in parallel with that of the ERP, with the P1-N170 sweep complex preceding the onset of fixation termination and subsequent image exploration, as indicated by the heatmap contrasts (Figure 10B). Thus, the time courses of the ERP and eye velocity show a clear relationship within individual trials and across participants in both tasks, face and affective picture viewing alike.

**Figure 10:**
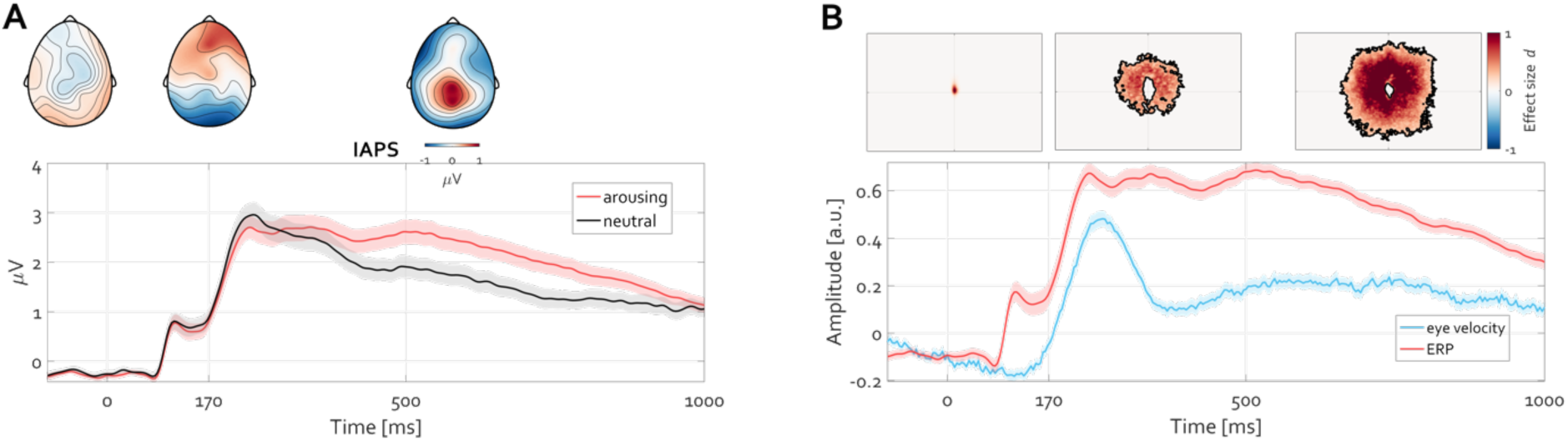
Grand average illustration of the initiation of eye movements following scene presentation. **A-** Grand-average ERP illustrating the initial P1-N170 sweep complex followed by a slow potential commonly referred to as the late positive potential (LPP- stronger for arousing compared to neutral images). The inset grand-average topographies illustrate the corresponding topography during the baseline, N170, and the difference (arousing minus neutral) for the LPP latency 400-1000 ms post-stimulus onset. **B-** On group level (N_group_ =85), eye movements toward image features follow after the P1-N170 ERP sweep complex. The line colors and design of the illustration are identical to Figure 9 but illustrate a group result rather than a single participant.

The temporal lag between the P1-N170 sweep complex and the onset of changes in eye velocity is nearly identical during face and picture viewing. However, it remains unclear to what extent the continued exploration behavior (e.g., sustained eye velocity and variability in gaze heatmap compared to fixation) unequivocally relates to the manifestation of slow cortical potentials. This aspect was evaluated next.

Under the hypothesis that condition differences in eye velocity are independent of differences in slow cortical potentials (commonly assumed but contrary to what this article proposes), the following consideration could be taken into account first. For each participant, the time course of eye velocity (e.g., 400 to 1000 ms) can be averaged to a single value per participant. This results in two distributions of eye velocity (one per condition, arousing and neutral), as depicted in Figure 11A, showing a significant condition difference (cluster permutation test, p<0.05). If one could eliminate these condition differences in eye velocity, then under the null hypothesis (advocated here) condition differences in slow cortical potentials (here, LPP) should remain unaffected. To evaluate this alternative, a procedure known as distribution stratification (https://www.fieldtriptoolbox.org/example/stratify/) can be applied. It involves removing values from non-overlapping parts of the distributions, thereby creating fully overlapping distributions and nullifying condition differences. Although this reduces participant numbers per condition, it prompts the question of whether eliminating eye velocity differences also eliminates LPP differences.

**Figure 11:**
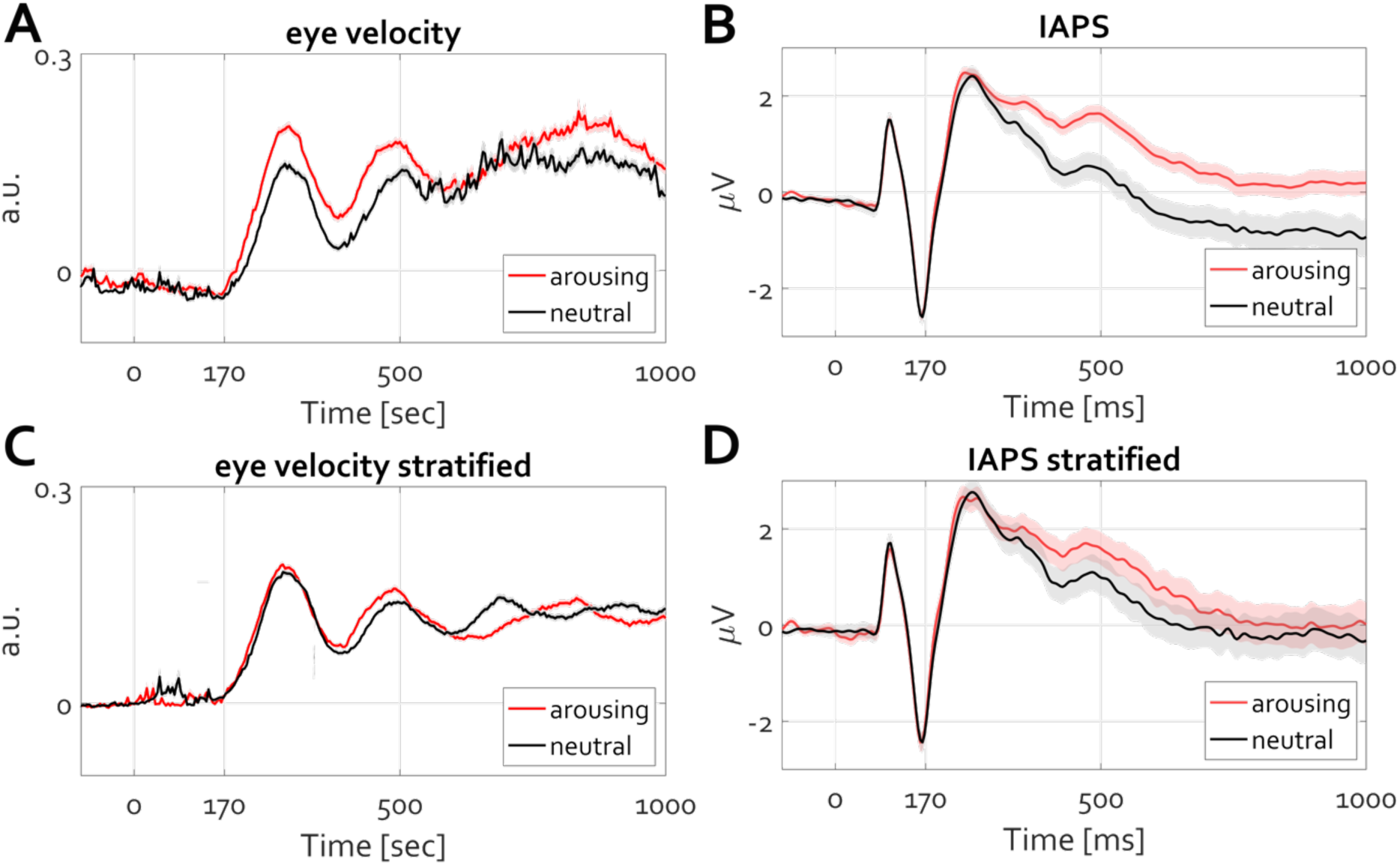
Eliminating condition differences within trials in eye velocity attenuates condition differences within trials in LPP. **A-** Eye velocity time course for the arousing (red line) and neutral (black line) conditions. The shading denotes the SEM across trials. A significant condition difference between eye velocity amplitude, larger for arousing than for neutral conditions, is evident. **B-** Similar to (A) but illustrating LPP. **C-** Eye velocity time course of stratified data highlighting no condition difference in eye velocity (e.g., means across the epoch do not differ). **D-** ERP average for the same trials in C demonstrating the absence of condition differences in LPP.

The increase in eye velocity following the P1-N170 sweep complex is notably stronger and more sustained for arousing scenes than for neutral scenes (Figure 11A, cluster permutation test, p < 0.05), as are the condition differences in slow cortical potential (Figure 11B, cluster permutation test, p < 0.05). Stratifying the distributions of eye velocity by averaging velocity values across the epoch and selecting trials with overlapping distributions diminish condition differences in eye velocity over time (Figure 11C). Although the procedure allows some of the latencies in the time courses to still exhibit significant condition differences, across the entire epochs the means do not differ, consistent with the null hypothesis. For the remaining trials, that is the stratified trials per condition for which there is no condition difference in eye movements, the ERP time courses are shown in Figure 11D. The effect size of condition differences in slow cortical potential is markedly reduced, with overlapping error values suggesting a potential challenge to falsify the null hypothesis typically in affective psychophysiology that emotional arousal does not affect cortical responses to affective stimuli. This conclusion contradicting the present view applies also to group data stratification to artificially generated images, which were neither validated nor rated for arousal and valence and were easily distinguishable by participants. Yet all AI images were easily spotted by the participants for obvious reasons apparent to interested readers (https://osf.io/q4mez/). The primary driver of the LPP effect in AI-generated images was the mere fact that termination of fixation is followed by eye exploration to evaluate the image (e.g., heatmaps in Figure 10B). In both cases—i.e., Figures 11 and 12—the loss of LPP difference supports the view that slow cortical potentials, such as the LPP, covary with increased eye exploration, as measured by the rise in eye velocity from the fixation baseline.

**Figure 12:**
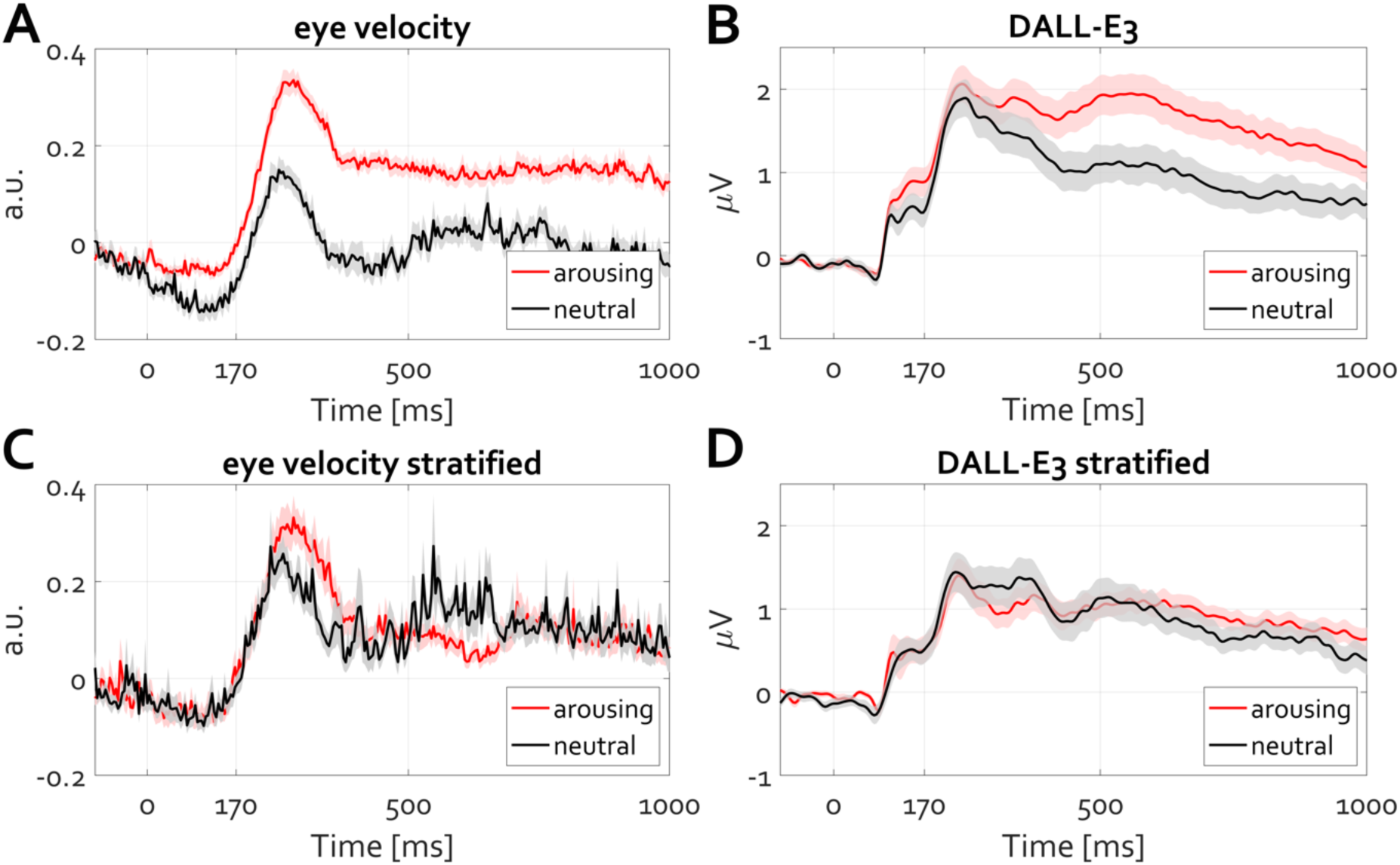
Eliminating condition differences within trials in eye velocity attenuates condition differences within trials in LPP. Stratification of the group-level (N=84) distributions for eye velocity eliminates group-level (N=25) condition differences in LPP. Figure design and outline identical to Figure 11. but for arousing images generated by artificial intelligence.

The informative value of analyzing time-frequency event-related modulation of ongoing oscillatory activity, although less frequently discussed than in the attention and working memory literature, has gained momentum in affective neuroscience (Codispoti, De Cesarei, & Ferrari, 2023; Flösch, Flaisch, Imhof, & Schupp, 2024; T. Popov, A. Steffen, N. Weisz, G. A. Miller, & B. Rockstroh, 2012; Schubring et al., 2020; Schubring & Schupp, 2019; Schubring & Schupp, 2020). The interpretation is that high-arousing stimulus material is associated with more reduction in alpha/beta power tan for neutral material, indicating cortical excitation and engagement of motivational circuits. Less clear, however, is how and why the reduction of alpha/beta power "reflects cortical excitability associated with the engagement of the motivational systems," as concluded in a recent review on alpha-band oscillations in emotion (Codispoti et al., 2023). Is this different from, for example, spatial or covert attention, and what does "excitability" actually excite? It cannot be the "motivational system" itself, as the term refers to a psychological construct in its purest form and by definition cannot be reduced to a constellation of excited neurons.

The present perspective suggests a complementary view. Similar to the contexts of face perception or previously reported spatial attention and working memory (Liu, Nobre, & van Ede, 2022, 2023; T. Popov, Gips, et al., 2023), as well as episodic memory formation (T. Popov & Staudigl, 2023), the reduction in alpha power aligns with the temporal occurrence of departure from fixation and ongoing image exploration (Figure 13).

**Figure 13:**
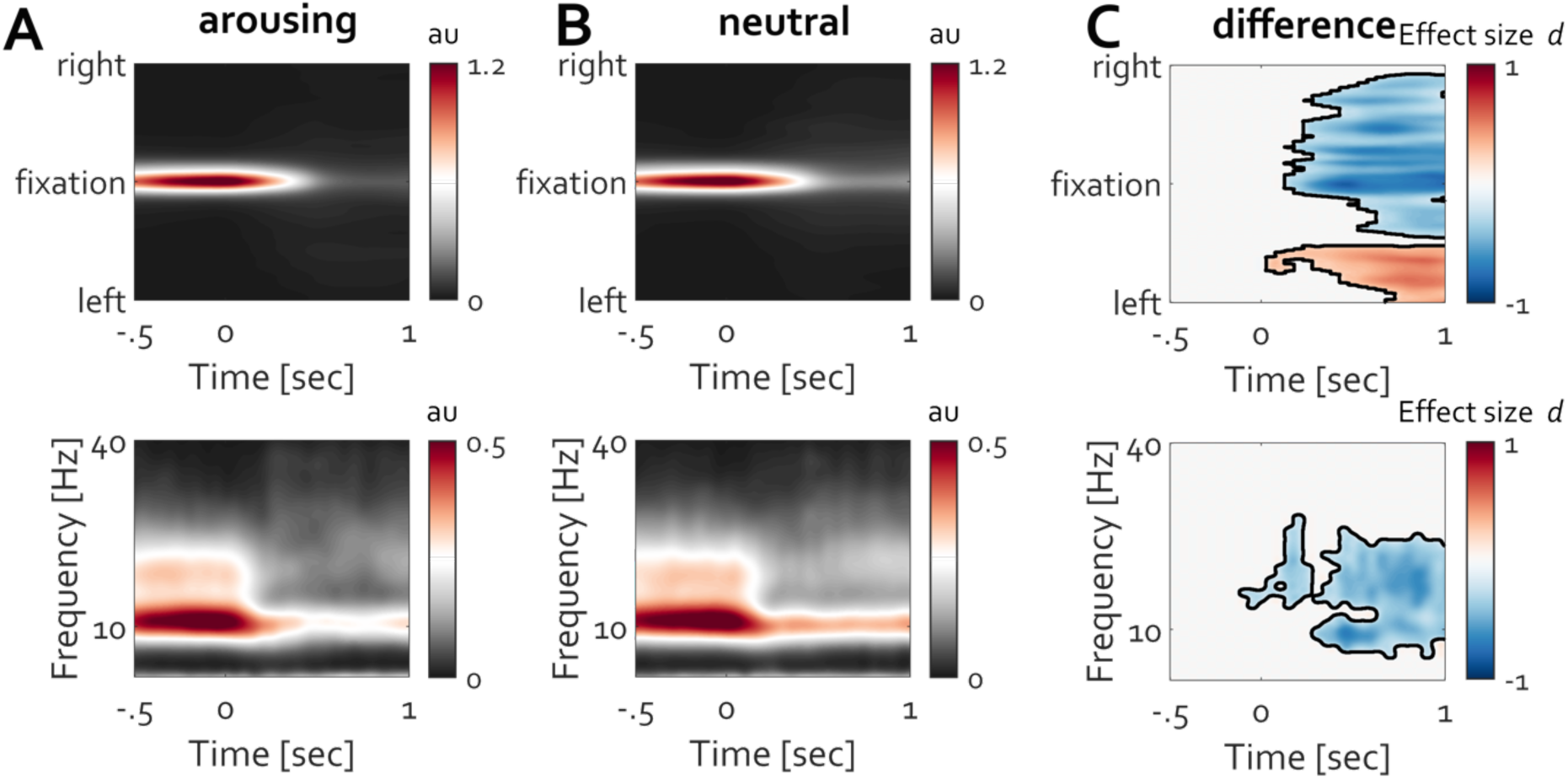
Occipital alpha power modulation and maintenance of fixation are related and inform departure from fixation during image exploration. **A-** Time-resolved histogram of gaze density around fixation (ordinate). The red color highlights the increased density of gaze positioning at fixation. Bottom- the corresponding time course of frequency-resolved activity averaged over electrodes O1 and O2. Warm colors indicate the strength of oscillatory power (in arbitrary units). The primary relevant finding is that the time course of power modulation corresponds to the time course of fixation maintenance. Departure from fixation toward exploring the face stimulus is associated with a reduction of alpha power. **B-** Identical to A but for neutral images from the IAPS database. **C-** Condition difference (arousing minus neutral) in gaze density (top) and alpha power (bottom). Confirmation of the typical reduction in alpha-beta power yet in association with more exploration for arousing than for neutral stimuli. Warm colors indicate an increase and cold colors a decrease in Cohen’s d effect size.

During the viewing of both arousing (Figure 13A) and neutral images (Figure 13B), fixation location is maintained during the prestimulus baseline. Subsequently, exploration of the image using eye movements is initiated, resulting in a decrease in occipital alpha power (Figure 13A, B, bottom). Neutral images are rated neutral not least due to their lower complexity, hence prompting less exploration by eye movements (Figure 13C, top, cluster permutation test, p <0.05). Consequently, the reduction in alpha power is stronger for complex images than for less complex neutral images (Figure 13C, bottom, cluster permutation test, p <0.05). This finding is nearly identical observation to that for covert attention and episodic memory formation (e.g., Figure 1B in (T. Popov & Staudigl, 2023)).

It is tempting to interpret this power modulation of alpha activity in association with eye exploration within the context of a motivational construct anchored along the approach-avoidance axis of behavioral disposition. Yet this association is ongoing, present throughout the awake state, as shown in the next analysis (publicly available data (Armeni et al., 2022)) of 10 hours of recordings of participants listening to audiobooks (Figure 14). Participants were tasked with listening to stories for 1 hour per session over 10 sessions during magnetoencephalography (MEG) and simultaneous eye tracking. (Participant-specific head casts enabled repositioning within the MEG helmet to minimize head positioning errors between sessions to acquire 10 hours of data.) At random intervals (from the participant’s perspective), written text appeared on the visual screen: questions about the narrative details and ensuring attention to the auditory narrative. Figure 14 illustrates the spectral energy of the MEG signal along with the corresponding time-resolved maintenance of gaze position at central fixation. Termination of the latter corresponds to the occurrence of story-related questions, necessitating consistent eye movement. Similar to the above results during face and image processing, the time course of variation in posterior alpha activity aligned with maintenance of and departure from fixation. This pattern is evident for all three available participants and was manifested near posterior-occipital sensors throughout the 10 hours (topographies in Figure 14, cluster permutation test, p < 0.05).

**Figure 14:**
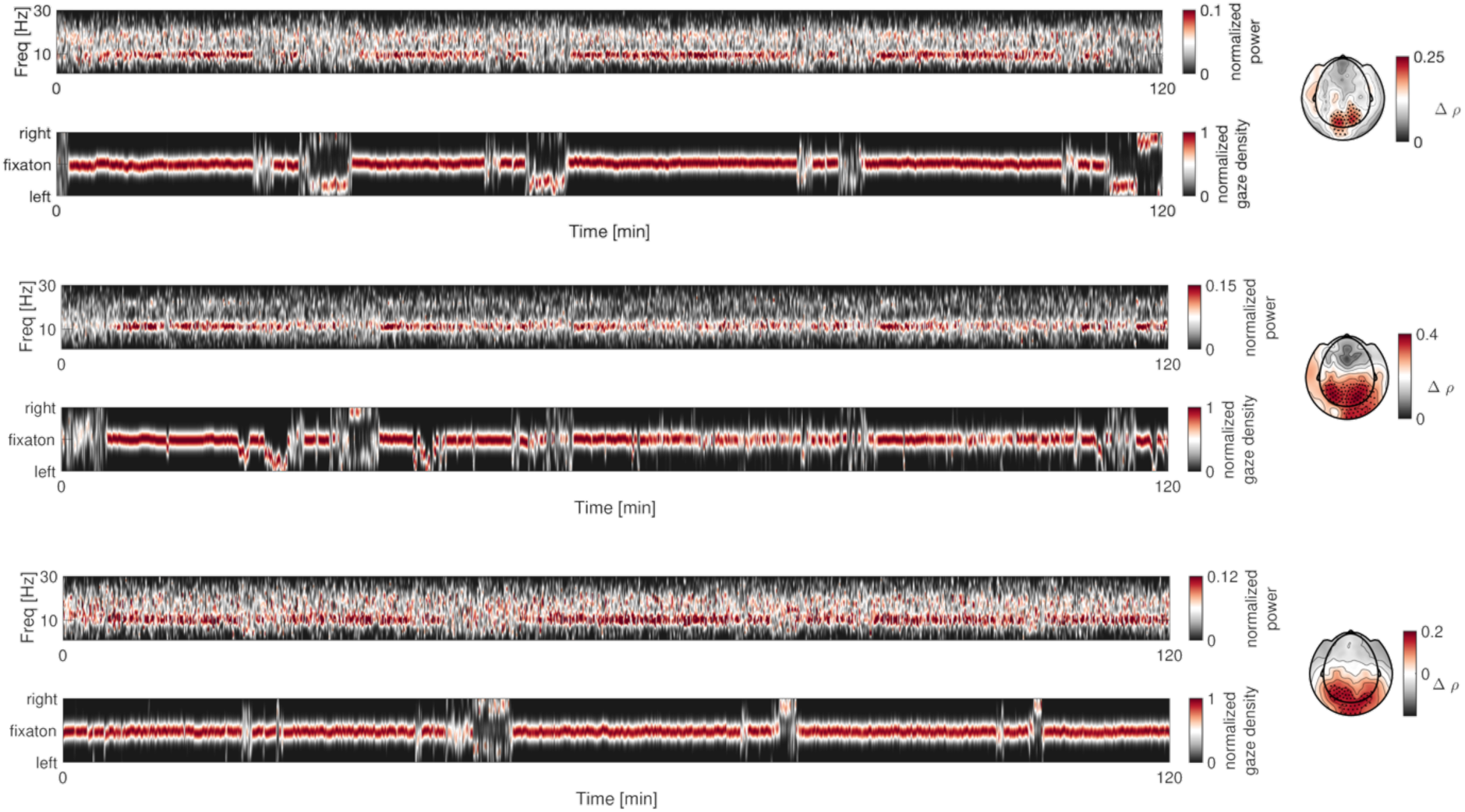
Maintaining gaze direction at fixation displays variations that parallel the modulation of alpha power in the visual cortex across a longer temporal scale. Two hours from a 10-hour dataset are illustrated for three participants. Their corresponding time-frequency spectra and fixation maintenance at the point of fixation are depicted, with the x-axis representing time and the y-axis representing frequency or fixation direction (up for rightward bias, down for leftward). The scalp topographies illustrate the correlation (cluster permutation test, p < 0.05) between alpha power and variability in gaze density throughout the entire 10 hours, contrasted against surrogate data obtained by circularly shifting the time series 500 times.

Several experiments narrated above support the present proposal that EEG components in common psychophysiological experiments relate to cognition indirectly through their more direct relationship with oculomotor action. However, relying solely on the motivational construct in affective picture viewing and face perception does not suffice to infer the asserted specificity of EEG-derived spectral measures in these contexts. Instead, ocular action and occipital alpha power modulation appear entangled, highlighting the intertwined nature of gaze direction control and occipital alpha oscillations (Tzvetan Popov et al., 2021). The necessity to move the eye in a particular direction along the text reveals the manifestation of this entanglement. Consequent upon this latter observation, reading direction (rightward as in English or leftward as in Farsi, Arabic, and Hebrew) should be dissociable thru both gaze direction and occipital alpha power, a hypothesis evaluated next.

Participants proficient in reading English (rightward, English skills sufficient to pursue studies at the local university) and Farsi (leftward, their native language) viewed 200 randomly presented words (100 per category, Figure 15A, top). The list of words can be accessed here (https://osf.io/q4mez/). Time- and time-frequency-domain analyses confirmed the prediction that reading direction entails consistent gaze bias and contralateral modulation of alpha oscillations and slow cortical potentials (Figure 15A, bottom). Also during reading, initiation of eye movement followed the P1-N170 ERP sweep complex, similar to what was observed for faces and complex images (e.g., Figures 5, 7, 9, 11).

**Figure 15:**
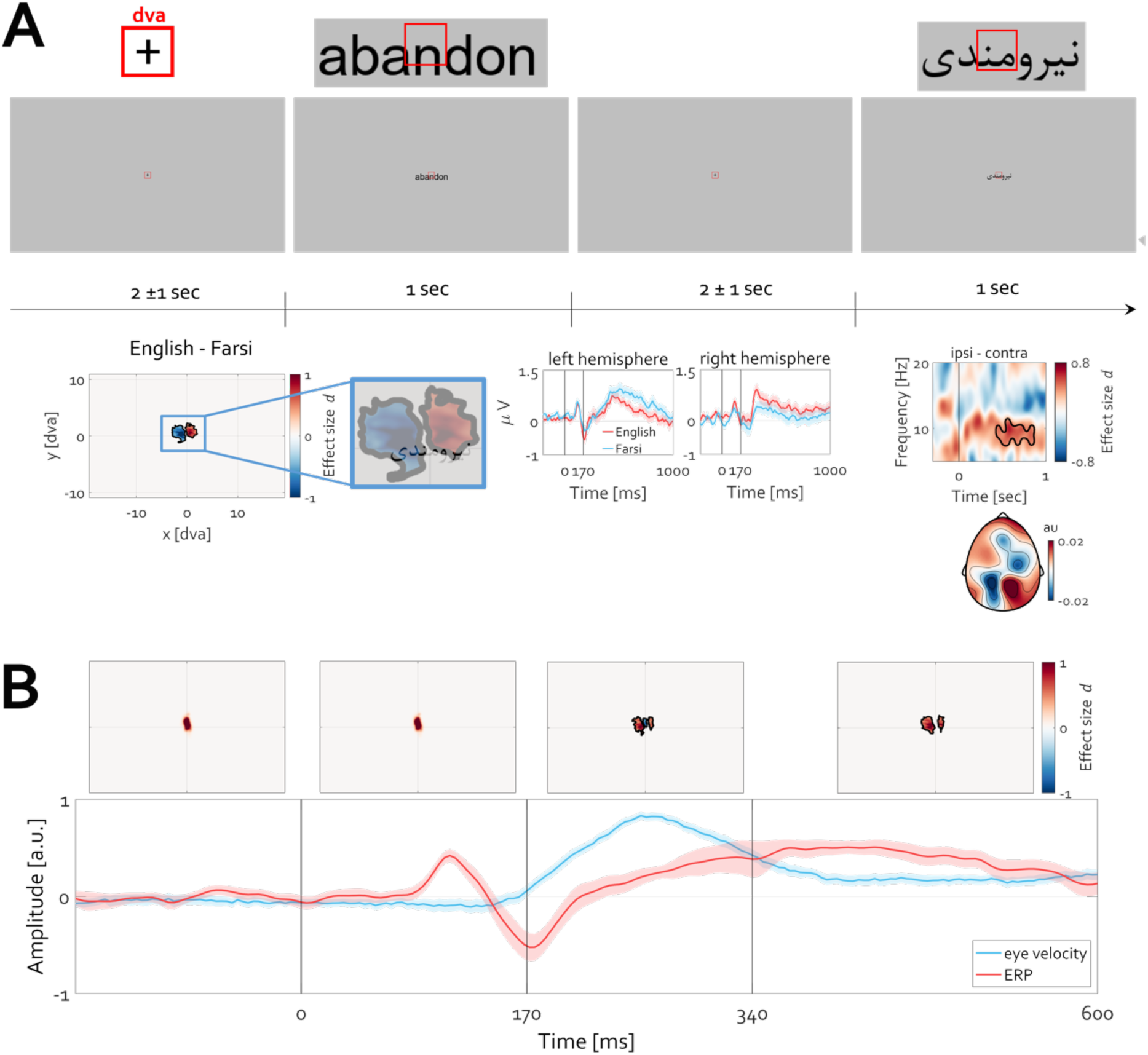
Slow brain potentials and alpha oscillations vary with reading direction. **A-**.Participants (N=28) capable of reading English and Farsi (mother tongue) words were tasked with silently reading words (pseudorandomly and centrally presented). Following the fixation period of 2±1 sec duration, words were presented for 1 sec. No task was involved except reading the words. Insets at the top illustrate examples of the size of the words relative to 1 dva. The time course of the experiment illustrated below the insets is at the true spatial viewing scale during acquisition. The contrast between the 2D histograms of gaze direction during English minus Farsi word reading revealed a clear lateralization of reading direction. The ERPs averaged across a set of left (T7 C3 P7 P3 M1) and right (T8 C4 P8 P4 M2) electrodes confirmed a clear difference in the amplitude of slow cortical potentials (cluster permutation test, p < 0.05). Alpha power was lateralized, with an occipital topography indicating that reading English words entails rightward eye gaze direction and contralateral left decrease in alpha power and vice versa. The contrast ipsi-minus contra-lateral to reading direction (O1 vs O2) revealed a significant condition difference illustrated in the time-frequency spectrogram (the black outline highlights a cluster of time-frequency activity supporting the rejection of the null hypothesis that alpha power does not depend on reading direction, cluster permutation test, p < 0.05. **B-** Initiation of eye movement required for reading follows after the P1-N170 ERP sweep complex. The time courses are identical to those reported in e.g. Figure 7. A statistically significant deviation in gaze position is observed after N170, indicating termination of fixation (blue colors in the heatmap denoting negative Cohen’s d effect size, cluster permutation test corrected for multiple comparisons, p<0.05) toward the letters in the word (red color in the heatmap denoting positive Cohen’s d effect size). English and Farsi conditions were pooled. The aggregated position of gaze for the time interval 0 to 170 ms is nearly identical to that extracted from the baseline (-170 to 0). The change in gaze position required for reading happens thereafter and coincides with the slow cortical potential in a way similar to that for faces, IAPS, and DALL-E3 images.

As a final line of evidence, how does the ubiquitous increase of alpha power during closed eyes relate to the present contention that EEG components in common psychophysiological experiments relate to cognition indirectly through their more direct relationship with oculomotor action? In 1932 Edgar Douglas Adrian reported the presence of alpha oscillations in both insect and human brains, which led him to initially refer to Hans Berger’s discovery 8 years prior as the Berger rhythm. However, Hans Berger declined that honor, and today it is universally known as the alpha oscillation (alpha as the highest observable amplitude, beta for the next highest, and so forth, under certain conditions). The most important conclusion drawn, which dominates current thinking, is that light and its absence are the primary reasons for the emergence of this rhythm as light off in the insect case and eyelid closure in the human case prompt increase of alpha oscillations amplitude. That interpretation dominates current understanding, resulting in other descriptions such as the “idling rhythm,” serving functional inhibition of cortical circuits and maintaining cerebral connectivity during eyes-closed rest. Aside the fact that these interpretations can be derived even in the absence of data, recent observations discuss the presence of a dominant alpha-like rhythm in the insect brain (honeybee) in relation to sensory action (similar to eye movement control through sensory antenna movement) (T. Popov & Szyszka, 2020). Amputation of the sensory antenna abolishes this rhythm (e.g. Figure 6d in (Tzvetan Popov et al., 2021)). That is, not light but action abolishes the rhythm; not darkness but inaction enhances the rhythm. What could be the mechanism of this effect? It is quite unlikely that eyelid closure per se modulates the amplitude of alpha oscillations.

Re-analysis of openly available data evaluating EEG during the eyes-closed state (T. Popov, Trondle, et al., 2023), as well as around microsleep events (Skorucak et al., 2019), suggests a profound difference despite the identical event (eyelid closure, Figure 16). Instructed closing the eyes increases occipital alpha oscillations (Figure 16A, averaged over electrode locations O1 and O2). The same event, eyelid closure, results in the opposite modulation around a microsleep event (Figure 16B), providing a complementary view of the control of the eyeball in orbit along the optic axis. During awake eyelid closure, the position is actively maintained under closed eyelids. From first-person experience, the struggle to maintain position along the optical axis before microsleep events is continuously increasing. Subsequent eyelid closure terminates this control, hence the reduction of the rhythm: a tractable empirical question.

**Figure 16:**
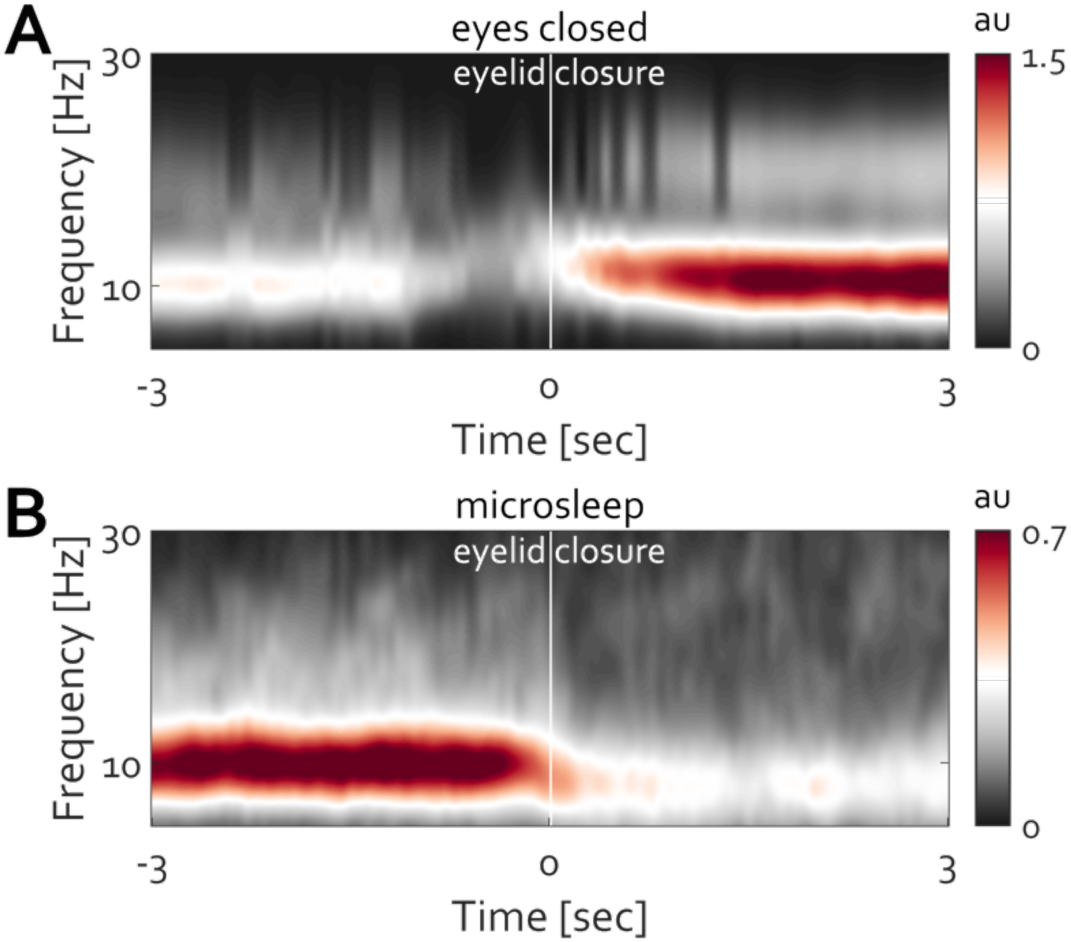
Eyelid closure results in distinct qualitative changes in alpha amplitude modulation. **A-** During wakefulness, closing the eyelids leads to an amplitude increase. **B**-In contrast, eyelid closure coinciding with the start of a microsleep event diminishes the amplitude of alpha activity.

## Discussion

The central premise of the present narrative is that the purpose of overt action is the control of perception. Crucially, concerning vision, the default state of the human eye is movement. Fixation intervals prior to stimulus presentation independent of the context and sensory modality studied are a common feature in nearly all psychophysiological experiments involving non-invasive electrical and magnetic neurophysiology. This task requirement in effect poses a difficult task for the human eye and requires control against its default state. The present report examined a variety of such experiments covering constructs such as social cognition, emotion, reading, and attention. Results foreground the association between both (a) ERP waveform components and power modulation of ongoing alpha oscillations and (b) the termination and the maintenance of fixation, respectively. As outlined above, the P1-N170 complex is used here to refer to what is typically known as the N170 in face processing literature and as the N1 in visual attention contexts including those in which emotion is probed. Traditionally, these different ERP component names imply distinct meanings. Using the term "P1-N170 complex" in the present work subsumes these terms, foregrounding the conclusion that, independent of the study context and its interpretation, the complex informs the experimenter about an upcoming oculomotor action triggered by the termination of fixation, rendering problematic the routine reliance on prestimulus baselines in psychophysiological research.

Termination of fixation and increase in eye velocity were found consistently after the manifestation of the P1-N170 ERP component across all tasks examined: passive viewing of faces, complex scenes, and reading. Slow sustained cortical potentials were associated with more (and more sustained) exploration of the presented face (e.g., Figures 4-7), visual scene (e.g., Figures 9-12), or word (Figure 15), highlighting their close relationship with ongoing oculomotor action enabling elaborate perception and evaluation. Furthermore, across all datasets presented here, maintenance of fixation was associated with occipital alpha power increase and termination thereof with alpha power reduction (e.g., Figures 8, 13, 14, 16). These EEG-derived metrics have been traditionally interpreted as the cortical manifestation of the cognitive construct studied, e.g. alpha power reduction as an indicator of covert attention in attention studies [e.g. (Foster & Awh, 2019; Foster, Sutterer, Serences, Vogel, & Awh, 2017; John J. Foxe & Adam C. Snyder, 2011; Ole Jensen, 2024; Klimesch, 2012)] or as an indication of engagement of motivational brain circuits in affective research [e.g. (Codispoti et al., 2023; Flösch et al., 2024; Tzvetan Popov, Astrid Steffen, Nathan Weisz, Gregory A. Miller, & Brigitte Rockstroh, 2012; Schubring & Schupp, 2019)]. The present contention is not that such interpretations are incorrect but that they depend on a largely ignored mechanism that mediates such relationships.

### Visual ERP Components Are Motor-Visual

The face N170 is arguably just a variant of N1 commonly in attention contexts. The N170 component is considered a manifestation of face processing in experiments utilizing face stimuli (Bentin, Allison, Puce, Perez, & McCarthy, 1996) but is considered an N1 (distinct in terminology and in functional significance) despite also occurring around 170 ms as an attention-selective component in relevant attention literature (Capilla et al., 2016; Luck, 1995; Vogel & Luck, 2000). Present results suggest a complementary view of these phenomena and their inferred meaning. The P1-N170 complex emerges following the instruction to maintain fixation where, upon stimulus presentation independent of the context (face, complex scene, or word), a motor-visual command is initiated, the result of which is an increase in eye velocity. It is the fixation instruction common to all of these tasks that provides a common explanation of why and how P1-N170 modulation is reliably observed across cognitive tasks and constructs studied.

The complementary view proposed here is that early visual ERP components prompt oculomotor action on which perception depends, revealing a motor function of the primary visual cortex in directing eye movements to a particular location in the environment (Peter H. Schiller & Edward J. Tehovnik, 2001; Schiller & Tehovnik, 2005). Consistent with this notion, the scalp topography of the P1-N170 complex is retinotopic. That is, it appears aligned with the order and direction in which light reflection hits the retina and is preserved along the visual pathway, eventually manifesting in interpretable EEG topography (Capilla et al., 2016; Dimigen & Ehinger, 2021; Kurki, Hyvärinen, & Henriksson, 2022; Woldorff et al., 1997). The general agreement is that neighboring objects stimulate neighboring retinal cells and manifest in neighboring voxels and or EEG electrodes. Such factors motivate the conclusion that hemodynamic or electromagnetic topographies in such a context confirm the retinotopic organization of the visual cortex (Capilla et al., 2016; Engel, Glover, & Wandell, 1997; Kastner, Pinsk, De Weerd, Desimone, & Ungerleider, 1999; Kurki et al., 2022; Wang, Mruczek, Arcaro, & Kastner, 2015). It follows from this that, if one removes the input to the retina, one would abolish the topography. But as mentioned above this does not happen. Instead, retinotopic topography is still present in full darkness (Tzvetan Popov et al., 2021), reminiscent of the report (Worden, Foxe, Wang, & Simpson, 2000) that sparked ongoing research detailing the functional significance of the retinotopic organization in spatial attention. That is, if the retinal drive is removed by being in a completely dark room, a retinotopic topography in the visual cortex is still present, as are hemifield-specific eye movements. In effect, the P1-N170 complex drives efferent oculomotor action. Accordingly, the ascribed specificity of the N170 component in encoding low-level face features is misleading.

And yet, if N170 happens (at least begins) too early for the fovea to have moved to see stimulus features that distinguish a face (e.g., Figures 4-7), how would one account for the general agreement that N170 is enhanced for face stimuli? A recent contribution of exceptional scholarship summarizes the past and current state of the art of eye-tracking research and breaks new ground in offering a comprehensive means of jointly analyzing electrophysiological and eye-tracking data to improve the evaluation of hypotheses in psychophysiological research (Dimigen & Ehinger, 2021). A finding of particular interest for the present narrative is the observation that P1-N170 amplitude is modulated by the size and direction of the saccades (e.g., Figure 7 in (Dimigen & Ehinger, 2021)). Specifically, when locked to fixation onset, the preceding saccade size and direction modulate the amplitude and retinotopic topography of the P1-N170 occurring within the first 300 ms after fixation onset. This fixation-related potential (FRP) is an electrophysiological phenomenon of its own, robust, replicable, and widely studied (Degno & Liversedge, 2020). However, a complementary interpretation of this FRP phenomenon can be offered. Humans typically saccade 3 to 6 times per second. It follows from this that fixation duration typically varies between approximately 150 and 300 ms (Tullis & Albert, 2013). Given an average fixation duration of 264 ms (per (Dimigen & Ehinger, 2021)), the occurrence of P1-N170s, rather than being evoked by the fixation, can also signify the cortical activity associated with the motor command preceding the upcoming saccade. That is, the duration between 0 ms (fixation onset) and 264 ms (average latency of subsequent saccade onset) includes the FRP. One can wonder whether this FRP is evoked by the fixation, as commonly accepted, or by the initiation of the subsequent eye movement that inevitably follows periods of fixation?

The present narrative argues for the latter. Amplitude variation with preceding saccade size as demonstrated by (Dimigen & Ehinger, 2021) does not necessarily support the assumption that this P1-N170 amplitude variation will also apply to an upcoming saccade. However, evidence also suggests that prior saccade direction influences the direction of future saccades (Anderson, Yadav, & Carpenter, 2008; Jones, Cowper-Smith, & Westwood, 2014). Specifically, this research shows that saccadic reaction times are reduced when the direction of a current saccade matches that of a preceding saccade, a phenomenon referred to as the "same direction benefit" (SDB) (Jones et al., 2014). This benefit persists even when intervening saccades occur between two same-direction saccades, indicating that saccade direction history influences future eye movements. Hence, it is conceivable that the retinotopic and amplitude modulation of P1-N170 also informs upcoming saccade direction akin to the retinotopic modulation of the saccadic spike potential (Keren, Yuval-Greenberg, & Deouell, 2010) and alpha oscillatory activity (T. Popov, Gips, et al., 2023; Tzvetan Popov et al., 2021; Quax, Dijkstra, van Staveren, Bosch, & van Gerven, 2019). In the case of faces, the pattern of eye movements follows the prototypical triangular pattern-left eye, right eye, mouth-repeat. As indicated in the heatmaps (e.g., Figures 4, 6, 7) and known since Yarbus (Yarbus, 2013), this pattern is well-established (Dimigen & Ehinger, 2021; Spiering & Dimigen, 2024). In the case of complex scenes, such as houses and IAPS images, the pattern of eye movements does not follow a consistent pattern; the saccades vary considerably in size and direction, without a clear, consistent spatial sequence across trials (e.g., Figures 9, 10). This is evident in the heatmaps for IAPS stimuli and when houses are used as stimulus material (Ward, 2017). That is, when viewing a house, eye movements averaged over viewers do not typically result in a pattern such as repeatedly fixating on the left roof, right roof, door, etc. It is thus conceivable that the amplitude of N170 is enhanced during face viewing due to the specific eye movements required. This proposal is supported by the established view that N170 is a robust phenomenon when controlling for stimulus complexity (e.g. (Johnston, Molyneux, & Young, 2014)). Yet, as argued above, this stimulus complexity involves large-amplitude saccades picturing the silhouette of a triangle when viewing faces vs. small-amplitude saccades and fixational eye movements when evaluating scenes.

Recent work concludes that it is saccade onset and not fixation onset as the behavioral event that triggers and best explains the manifestation of early visual evoked responses such as P1-N170 discussed here (Amme et al., 2024). This proposal contradicts the FRP literature and challenges the notion that a fixation event by itself can elicit P1-N170. Instead, Amme and colleagues noted that saccade duration and therefore amplitude (as described by the saccade “main sequence” (Bahill, Clark, & Stark, 1975)) predict the peak latency of the P1 component. When locked to fixation onset, the authors observed that the shorter the saccade, the longer the P1 peak latency, and vice versa. In addition, small saccades were associated with smaller P1 amplitude and vice versa. That is, the amplitude and duration of the oculomotor event are closely linked to the amplitude and latency of P1-N170.

Accepting one premise of the present proposal, that the default state of the eye is movement, and rejecting the assumption of fixation as a stationary event independent of continuous, active oculomotor control, these seemingly contradictory findings can be reconciled by considering the possibility that the strategy of locking the data on fixation or saccade onset provides a view of the same oculomotor event from two different vantage points, thus both valid observations representing the variability in the electrophysiological data at the rise of P1-N170 when locked to fixation onset and at the decline when locked to saccade onset. In fact, a closer look at both Dimingen and Ehinger arguing for FRP (e.g., Figure 7F in (Dimigen & Ehinger, 2021)) and Amme and colleagues arguing against it (e.g., Figure 1D in (Amme et al., 2024)) reveals fair agreement in that in both reports – saccade size prior to fixation onset (Dimigen & Ehinger, 2021) or saccade size following the fixation onset (Amme et al., 2024) – the same phenomenon is observed: the smaller the saccade amplitude, the later the P1-N170 peak latencies and the smaller their peak amplitudes. The converse appears also true. Consideration of the timing and conduction delay starting at the level of the retina, lateral geniculate nucleus (LGN) in the thalamus, and the early entry into V1 gives some hints as to why this is the case.

Recent work utilizing state-of-the-art, simultaneous recordings of retinal potentials (electroretinogram, ERG) and MEG provides the most direct clues to date in a human model system (Britta U. Westner & Dalal, 2019; B. U. Westner, Lubell, Jensen, Hokland, & Dalal, 2021). The authors measured two retinal responses, the *a-wave* and *b-wave*, which are ERG components that reflect retinal activity in response to visual stimuli. These waves reflect specific stages of retinal signal processing before the information reaches higher visual areas such as the LGN and V1. The a-wave that peaks at ∼25ms represents the initial hyperpolarization of photoreceptor cells (rods and cones) in response to light, thus very early stages of retinal signal transduction, converting light into electrical signals. The b-wave that peaks at ∼80ms reflects inner retinal activity, primarily from bipolar cells, which process and relay the photoreceptor signals to ganglion cells – a critical stage of signal integration and amplification within the retina. In fact, this is the timing of arrival of the retinal signal in the primary visual cortex reported non-invasively in humans (Britta U. Westner & Dalal, 2019). The latest time point at which V1 was informed about the events occurring on the retina was 77 ms. That is, from this point on, the pyramidal cells in V1 are capable of sending a signal down to SC, the brain stem, the eye muscle, and eventually directly the eye in motion toward a particular position at the visual display. As mentioned in the introduction, this is the chain of events observed in the non-human primate brain that is described in detail elsewhere (Schiller et al., 1974; Schiller & Tehovnik, 2003). Important here is that saccade initiation by the SC depends on the motor command from V1, i.e. a saccade initiation to terminate the experimentally defined state of fixation maintenance and explore a face, complex image, word, etc. Thus, after ∼77 ms at the latest, this command is issued downstream along the proposed circuit of saccade generation (Tehovnik, Slocum, Carvey, & Schiller, 2005) - to SC, brainstem nuclei, and eye muscles. The duration of the transmission is at least ∼100 ms until an eye tracker records a movement of the eye in orbit [e.g.(Fischer & Ramsperger, 1984; Schiller, Haushofer, & Kendall, 2004; Stine et al., 2023)]. Summing these numbers, one derives a latency of ∼177 ms, prior to which eye movement following stimulus onset appears unlikely. This estimate is in line with present results confirming the observation that, at least in part, the functionality of early evoked responses, besides being visual/perceptive, is also motor – directing the mind’s eye (attention) and the observer’s eye (sensor) toward a state of active exploration of the environment. Put simply, observer-generated action is in charge of perception.

### Amplitude Modulation of the Late Positive Potential (LPP) Varies with the Intensity of Exploration via Oculomotor Action

To reiterate, the purpose of overt action/behavior is the control of perception. According to the bio-informational theory of emotional imagery (Lang, 1979), the perceptual response itself results from an efferent motor program as a prototype for overt behavior. The theory was a foundational view instrumental in the development of databases such as the International Affective Picture System (IAPS) for the systematic study of emotion, including but not limited to EEG. Consequently, the LPP became among the more widely studied phenomena in affective psychophysiology. It is considered an important electrophysiological manifestation of affective stimulus processing. Present results suggest a mechanism for how and why the LPP is a robust, replicable metric, the enhancement of which is widely considered to reflect “selective processing of emotional stimuli” (Cuthbert et al., 2000).

Here is how the present theory explains the LPP effect. In order to evaluate the stimulus, emotional stimuli motivate oculomotor exploration [in accord with (Lang, 1979)]. Eliminating the difference in oculomotor exploration diminishes the condition difference in LPP amplitudes both on individual (Figure 11) and group levels (Figure 12). That is, the specific electrophysiological response triggered by the very same high-arousing images is attenuated in the absence of condition differences in eye movements to the extent that the initially rejected null hypothesis is no longer falsifiable (e.g., Figure 11D, Figure 12). Furthermore, research often utilizes well-standardized photos of real-world settings, such as the IAPS database, for obvious reasons. However, the presence of a nearly identical LPP effect triggered by artificial images – despite these images not being rated or validated to ensure their construct validity, and being clearly identified by participants as not real – requires interpretation. This interpretation aligns with the current state of the art, which foregrounds the preservation and automatic utilization of neural circuits signaling threat and danger in all cases, whether real or artificial, in which the propositional structure of threat or danger are initiated (Lang, 1979). Thus, LPP enhancement or attenuation to IAPS and AI images is likely of similar origin. The present work is in line with such interpretation yet foregrounds the fact that, in both cases, images are evaluated by exploratory eye movements, the attenuation of which attenuates the LPP effect in both cases – standardized images of real-life scenarios and artificial images. Hence, the present complementary view is that LPP responses indeed relate to emotion but via their primary relation to eye exploration. Ongoing eye exploration is not specific to affective images but is also present in the case of faces and word reading where they also co-vary with slow cortical potentials (e.g., Figures 5, 6, and 15). In support of this conclusion, Kurki and colleagues provide compelling evidence of retinotopy modulation in slow sustained cortical fields (Kurki et al., 2022), mirroring findings in the context of reading presented here. The critical observation is: whether the reading process necessitates eye movements to the left or right, or whether an attention probe in the left or right visual field triggers corresponding eye movements, the phenomenon remains consistent. In both scenarios, slow cortical potentials are observed to align with the direction of oculomotor action.

### Quo vadis Alpha Oscillations?

The frequency of reports highlighting a relationship between the direction and variability of eye movements and power modulation of alpha oscillations is increasing (Balestrieri, Michel, & Busch, 2024; Cruz et al., 2024; Kornrumpf, Dimigen, & Sommer, 2017; Liu et al., 2022, 2023; T. Popov, Gips, et al., 2023; Tzvetan Popov et al., 2021; T. Popov & Staudigl, 2023). These studies consistently report a bias in the direction and variability of microsaccades contralateral to the scalp-manifested modulation of alpha power. Interpretations and conclusions vary within the scope and boundaries of the construct studied in a given study. Yet, at minimum, the directional bias of microsaccades appears entangled with contralateral alpha power modulation. That is, either informs the state of the other. Based on the results presented here, this relationship appears continuous and ongoing: reduction in eye movement to maintain fixation prompts high alpha amplitudes and vice versa. Alpha power reduction is always associated with an increase in eye movements over baseline. This relationship is observed in all cases studied here, independent of task and context, and is evident on different temporal scales, seconds (e.g., Figure 8,13), minutes [e.g., Figure 3 in (T. Popov & Staudigl, 2023)], and hours (e.g., Figure 14). What is going on?

As mentioned in the introduction, maintaining fixation requires motor control, which can only be orchestrated by the brain. Hence, it is worth speculating about the signals and mechanisms that enable this control.

The present narrative argues that alpha oscillations play a critical role in this process. As mentioned earlier, neurons within the layered structures of the SC, V1, and parietal cortex are equipped with vector-coding firing capability. This means that the activation of a neuron associated with a specific position will always direct the eye to that position, irrespective of the initial starting point. Consequently, all possible positions are encoded by different neurons in each of these structures, enabling the full degrees of freedom required to explore the environment (within the constraints of morphology, field of view, etc.). Theoretically, when vectors are uniformly distributed across a circular space, each direction is equally likely. For every vector pointing in one direction, there is an equal and opposite vector pointing in the opposite direction. The vector sum of all such pairs cancels out. Given the premise of this narrative – that the default state of the eye is movement – it follows that no single neuron exists solely to maintain fixation at rest, but instead to program a specific direction of ocular movement [as demonstrated for example by (Schiller et al., 1974; P. H. Schiller & E. J. Tehovnik, 2001)]. Yet mammals do fixate, utilizing small fixational eye movements around a given target. If the neurons directing eye movement toward particular directions are uniformly distributed across the visual field and contribute equally to the net vector, their summed activity would cancel out, resulting in a zero vector. This implies no net movement, which could correspond to a neutral, forward-facing gaze along the optic axis. The present contention is that a key part of the mechanism by which this neuronal recruitment is achieved in the brain is alpha oscillations and their amplitude variation observed in all evaluations examined to date. The amplitude variation and the associated phase of opportunity for neuronal discharge have been reported and discussed in the context of alpha oscillations several times (Chapeton, Haque, Wittig, Inati, & Zaghloul, 2019; Haegens, Nácher, Luna, Romo, & Jensen, 2011; Huang et al., 2021; T. Popov & Szyszka, 2020; Saalmann, Pinsk, Wang, Li, & Kastner, 2012). Thus, based on present results one can speculate that an increase in alpha activity, with considerable volume conduction over occipital and parietal cortices, is an emergent event arising from the collective agency of neurons sensitive to different visual field locations, with the phase of the emergent rhythm exceeding the SNR level to an extent that in turn biases their firing properties. The consequence of this is the alignment of gaze along the optic axis. As the occurrence of behaviorally relevant events within the visual field is theoretically unpredictable, in the sense that any location within the visual field is equally possible, positioning the eye along the optic axis provides the best possible subsequent reaction and accuracy in movement toward an upcoming event; that is, it has the most of degrees of freedom.

Present data (Figures 8, 13, 14, 16) (and, I predict, all existing data utilizing fixation during baseline) support this conjecture: periods of fixation during the baseline are always associated with increased amplitudes of alpha activity due to the fixation requirement and decreased during subsequent eye exploration. I am aware of no exception in healthy human non-invasive psychophysiology. In fact, pioneering work that sparked subsequent research on the role of pre-, peri- and post-stimulus alpha in attention has already reported this: “We found significantly larger parieto-occipital ∼10 Hz activity in the period preceding a compound audio-visual stimulus when subjects were cued to selectively attend to the upcoming auditory features and ignore the concurrent visual features.”[p. 3932 in (Foxe, Simpson, & Ahlfors, 1998)]. The then and still prevailing interpretation is that the parieto-occipital cortex is actively engaged in anticipation. But how so? By initiating oculomotor action – that is the mechanism that has been missing in this literature. Thus, a plausible interpretation so far not considered is that, when participants anticipate a sound, maintenance of fixation is better achieved when they prepare to engage in a visual task. This would explain the difference in pre-stimulus alpha power, which, in addition, demonstrated already the increase in slow cortical potential in anticipation of visual demand-hence more eye movements.

It follows from this argument that the most compelling case of an increase in alpha oscillations, under closed eye lids, should actually maintain the eye position along the optical axis. This is in fact what we observe when we close eye lids: upon opening, the empirical sense is that our gaze is always straight ahead. Something, however, has to maintain this position to end up straight ahead as we open the eye lids, and this something is the alpha oscillations. In an awake vigilant state, eyelid closure prompts an increase in alpha power with a parieto-occipital scalp topography (Adrian & Matthews, 1934). From first-person experience, the eyes and gaze direction are aligned along the optic axis upon eyelid opening, but the mechanisms by which eye position under closed eyelids is maintained are a matter of ongoing research (Allik, Rauk, & Luuk, 1981; Ben Barak-Dror et al., 2024; Bergamin, Bizzarri, & Straumann, 2002; Iwasaki et al., 2005; Kirchner, Watson, Bauer, & Lappe, 2022). The general agreement is related to Bell’s phenomenon, a reflex movement associated with a slight upward and outward movement of the eyes that occurs when the eyelids are closed. The present narrative and results presented in Figure 16 support the conjecture that alpha power increase during closed eyelids supports the maintenance of eye position along the orbital axis, ensuring the readiness to act and see upon eyelids opening – that is, starting from the best possible position with a maximum degree of freedom, given the uncertainty of the upcoming optic flow. This starting position is straight ahead for the same reason detailed above. Conversely, struggling to maintain gaze direction along the optical axis, as is the case in situations leading up to microsleep events, the alpha power increase is observed before the eyelid closure and not after it, as the latter no longer entails active gaze control maintenance (Figure 16). In short, the three qualitatively different states of fixation maintenance (during baseline periods of cognitive tasks, maintenance of eye position along the optic axis during awake closed eyelid state, and before microsleep events) are supported by the same cortical phenomenon: an increase in alpha power temporally reduces the degrees of freedom of gaze-direction-selective neuronal firing in the oculomotor system.

The current debate in the literature about the role of alpha oscillations foregrounds the inconsistency in the empirical evidence as to whether and how modulations of alpha activity aid distractor suppression (Foster & Awh, 2019; Ole Jensen, 2024; Redding & Fiebelkorn, 2023). At the core of the argument is the limited evidence of a genuine increase in alpha power actively suppressing processing of distracting stimuli (Bonnefond & Jensen, 2012; J. J. Foxe & A. C. Snyder, 2011; O. Jensen & Mazaheri, 2010). Instead, the decrease in amplitude and the varying levels of it may support stimulus selection through signal enhancement (Foster & Awh, 2019). The most recent view is that power modulation of alpha activity is controlled by an indirect mechanism that regulates “the load of goal-driven information that promotes the alpha power in regions processing potentially distracting task-irrelevant information” (p. 6-7 in (Ole Jensen, 2024)). Consequently, alpha power modulation can but is not necessary under direct top-down control. That would explain the inconsistency in the empirical evidence for the distractor account of alpha oscillations (Ole Jensen, 2024), which in turn motivates the future research direction to “uncover which regions and neural mechanisms are involved in the indirect control and generation of alpha oscillations” (p.7 in (Ole Jensen, 2024)).

The present position that action controls perception provides a parsimonious explanation that does not require homuncular accounts (e.g., a top-down controller). Perceptual load is actively controlled by oculomotor action, with maintenance of fixation reducing load. That oculomotor action entails alpha power increase, whereas active exploration (i.e., looking) fosters perception (i.e., seeing). The regions involved, i.e. FEF, V1, parietal cortex, and subcortical areas, not only are similar within research detailing the cerebral mechanisms of oculomotor control (Peter H. Schiller & Edward J. Tehovnik, 2001) and research discussing circuits involved in alpha oscillation generation and maintenance (Bollimunta, Chen, Schroeder, & Ding, 2008; Halgren et al., 2019; Ole Jensen, 2024; Saalmann et al., 2012) but likely serve the same function: controlling action in the service of regulating perceptual load.

### Consequences of Misinterpreting Electrophysiology in Human Cognitive Neuroscience

The covariance of oculomotor action with cortical activation in psychophysiology has been noted repeatedly [e.g. (Carl, Açık, König, Engel, & Hipp, 2012; Dimigen, Valsecchi, Sommer, & Kliegl, 2009; Liu et al., 2023; Meyberg, Werkle-Bergner, Sommer, & Dimigen, 2015; Yuval-Greenberg, Tomer, Keren, Nelken, & Deouell, 2008)] including in one of the first reports on visual evoked responses (e.g., Figure 2 in (Chapman & Bragdon, 1964)). Recording the ERG, EOG, and EEG simultaneously, the 1964 paper concluded that the P1-N170 complex (not referred to as such at that time) was due to the cognitive operation under study rather than to oculomotor action. The latter was found to occur *after*, and not simultaneously with, the ERP complex and was therefore deemed irrelevant to the interpretation of the measured ERP.

This conclusion has been widely accepted in psychophysiological research the past 60 years, manifesting in the predominant premise that ERP components reflect the cognitive operations under study. However, as the present narrative argues, this conclusion can be revisited by asking: how so? Oculomotor action is not merely an artifact that can be traditionally addressed through data preprocessing and statistical removal. Instead, it is itself “a genuine cortical activity and not an EEG-specific artifact … also frequently overlaid on magnetoencephalographic and possibly hemodynamic datasets” [p. 12330 in (Dimigen et al., 2009)]. Therefore, as outlined above, misinterpretations and conclusions derived from EEG and MEG data encourage neglect of complementary alternatives and result in misjudgments with potentially serious consequences.

“*The dominant discourse in modern cognitive, affective, and clinical neuroscience assumes that we know how psychology/biology causation works when we do not … there are serious intellectual, clinical, and policy costs to pretending we do know … crucial scientific and clinical progress will be stymied as long as we frame psychology, biology, and their relationship in currently dominant ways*.”[p. 716 in (Miller, 2010)].

In this piece Miller argues for the illuminating power of studying biological phenomena in understanding psychological phenomena but argues against the reducibility of the latter to the former. The piece foregrounds and calls for more awareness of possible misinterpretation of their relationship(s) to reduce serious consequences for public policies and clinical practice implications.

And yet misinterpretation can be a contributor to scientific advance (e.g., earth-centric vs. heliocentric universe), unless considered an undoubted, proven fact, whereas Popperian logic forbids the notion of “proof” and foregrounds plausibility as the sole arbitrator. One important implication of the present perspective is that developing a chemical that will target modulation of the latency and/or amplitude of N170 in children with autism spectrum disorder (in hopes, for example, of improving their attention to and interpretation of others’ facial expressions) will potentially *also* impact their reading development (e.g., Figure 15), exploration of the environment (e.g., Figures 9, 13), and social cognition skills (e.g., Figures 5, 7, 8). Common rationales for, interpretations of, and reliance on results of drug development and clinical practice/treatment efficacy studies (Kala et al., 2021) are at best premature as bases for routine clinical intervention because the assumed scope of their effects is often too narrow. Evaluating effects just on target symptoms and assessing unwanted side effects sample too little of the potential impact space of such interventions. A broader conceptualization of the relevant mechanisms is needed.

In closing, I invite the reader to consider the following synthetic experiment. Let us suppose that there are only five neurons in primary motor cortex, each controlling the full degrees of freedom in the motion of the five hand digits. In a Braille reading experiment, words describing living vs. non-living objects are presented. State-of-the-art analysis will likely affirm that the recorded neural firing pattern can be used to decode word meaning. This conclusion seems obvious and likely. However, the neurons and therefore their activity pattern cannot and do not contain “meaning,” as their sole design is to move the respective digits. Meaning requires a trained observer and is an emergent property of that observer, not merely “information” contained in the recorded neural pattern. What is contained in the neural pattern is the plan of action, such as the control of the movement over the Braille pattern. Similarly, ERP waveforms and EEG oscillations in human cognitive neuroscience may be used to infer shape, color, facial features, or hedonic content. I submit the complementary view that ERP waveforms and oscillations in non-invasive visual electrophysiology (and likely in other contexts) primarily signify oculomotor action in the service of control of actions, the result of which is perception.

## Acknowledgements

The author would like to express his deepest gratitude to Gregory A. Miller for his guidance, insightful discussions, and contributions to earlier versions of this manuscript.

The author would like to thank Tobias Flaisch and Harald Schupp for lending some of the EEG hardware and providing the IAPS IDs used in one of the studies. The author also extends gratitude to all the participants who volunteered for the experiment.

## References

Adrian, E. D., & Matthews, B. H. C. (1934). THE BERGER RHYTHM: POTENTIAL CHANGES FROM THE OCCIPITAL LOBES IN MAN. Brain, 57, 355–385.

Allik, J., Rauk, M., & Luuk, A. (1981). Control and sense of eye movement behind closed eyelids. Perception, 10(1), 39–51. doi:10.1068/p100039

Amme, C., Sulewski, P., Spaak, E., Hebart, M. N., König, P., & Kietzmann, T. C. (2024). Saccade onset, not fixation onset, best explains early responses across the human visual cortex during naturalistic vision. bioRxiv, 2024.2010.2025.620167. doi:10.1101/2024.10.25.620167

Anderson, A. J., Yadav, H., & Carpenter, R. H. (2008). Directional prediction by the saccadic system. Curr Biol, 18(8), 614–618. doi:10.1016/j.cub.2008.03.057

Armeni, K., Güçlü, U., van Gerven, M., & Schoffelen, J. M. (2022). A 10-hour within-participant magnetoencephalography narrative dataset to test models of language comprehension. Sci Data, 9(1), 278. doi:10.1038/s41597-022-01382-7

Bahill, A. T., Clark, M. R., & Stark, L. (1975). The main sequence, a tool for studying human eye movements. Mathematical Biosciences, 24(3), 191–204. 10.1016/0025-5564(75)90075-9

Balestrieri, E., Michel, R., & Busch, N. A. (2024). Alpha-Band Lateralization and Microsaccades Elicited by Exogenous Cues Do Not Track Attentional Orienting. eneuro, 11(2), ENEURO.0076-0023.2023. doi:10.1523/eneuro.0076-23.2023

Ben Barak-Dror, O., Hadad, B., Barhum, H., Haggiag, D., Tepper, M., Gannot, I., & Nir, Y. (2024). Touchless short-wave infrared imaging for dynamic rapid pupillometry and gaze estimation in closed eyes. Communications Medicine, 4(1), 157. doi:10.1038/s43856-024-00572-1

Bentin, S., Allison, T., Puce, A., Perez, E., & McCarthy, G. (1996). Electrophysiological Studies of Face Perception in Humans. J Cogn Neurosci, 8(6), 551–565. doi:10.1162/jocn.1996.8.6.551

Bergamin, O., Bizzarri, S., & Straumann, D. (2002). Ocular torsion during voluntary blinks in humans. Invest Ophthalmol Vis Sci, 43(11), 3438–3443.

Bollimunta, A., Chen, Y., Schroeder, C. E., & Ding, M. (2008). Neuronal mechanisms of cortical alpha oscillations in awake-behaving macaques. J Neurosci, 28(40), 9976–9988. doi:10.1523/jneurosci.2699-08.2008

Bonnefond, M., & Jensen, O. (2012). Alpha Oscillations Serve to Protect Working Memory Maintenance against Anticipated Distracters. Current Biology, 22(20), 1969–1974. doi:10.1016/j.cub.2012.08.029

Capilla, A., Melcón, M., Kessel, D., Calderón, R., Pazo-Álvarez, P., & Carretié, L. (2016). Retinotopic mapping of visual event-related potentials. Biological Psychology, 118, 114–125. 10.1016/j.biopsycho.2016.05.009

Carl, C., Açık, A., König, P., Engel, A. K., & Hipp, J. F. (2012). The saccadic spike artifact in MEG. Neuroimage, 59(2), 1657–1667. 10.1016/j.neuroimage.2011.09.020

Chapeton, J. I., Haque, R., Wittig, J. H., Jr., Inati, S. K., & Zaghloul, K. A. (2019). Large-Scale Communication in the Human Brain Is Rhythmically Modulated through Alpha Coherence. Curr Biol, 29(17), 2801–2811.e2805. doi:10.1016/j.cub.2019.07.014

Chapman, R. M., & Bragdon, H. R. (1964). Evoked Responses to Numerical and Non-Numerical Visual Stimuli while Problem Solving. Nature, 203(4950), 1155–1157. doi:10.1038/2031155a0

Codispoti, M., De Cesarei, A., & Ferrari, V. (2023). Alpha-band oscillations and emotion: A review of studies on picture perception. Psychophysiology, 60(12), e14438. doi:10.1111/psyp.14438

Cruz, G., Melcón, M., Sutandi, L., Palva, J. M., Palva, S., & Thut, G. (2024). Oscillatory brain activity in the canonical alpha-band conceals distinct mechanisms in attention. *The Journal of Neuroscience*, e0918242024. doi:10.1523/jneurosci.0918-24.2024

Cuthbert, B. N., Schupp, H. T., Bradley, M. M., Birbaumer, N., & Lang, P. J. (2000). Brain potentials in affective picture processing: covariation with autonomic arousal and affective report. Biol Psychol, 52(2), 95–111. doi:10.1016/s0301-0511(99)00044-7

Degno, F., & Liversedge, S. P. (2020). Eye Movements and Fixation-Related Potentials in Reading: A Review. Vision (Basel), 4(1). doi:10.3390/vision4010011

Dimigen, O., & Ehinger, B. V. (2021). Regression-based analysis of combined EEG and eye-tracking data: Theory and applications. J Vis, 21(1), 3. doi:10.1167/jov.21.1.3

Dimigen, O., Valsecchi, M., Sommer, W., & Kliegl, R. (2009). Human microsaccade-related visual brain responses. J Neurosci, 29(39), 12321–12331. doi:10.1523/jneurosci.0911-09.2009

Donoghue, T., Haller, M., Peterson, E. J., Varma, P., Sebastian, P., Gao, R., Voytek, B. (2020). Parameterizing neural power spectra into periodic and aperiodic components. Nature Neuroscience, 23(12), 1655–1665. doi:10.1038/s41593-020-00744-x

Engel, S. A., Glover, G. H., & Wandell, B. A. (1997). Retinotopic organization in human visual cortex and the spatial precision of functional MRI. Cereb Cortex, 7(2), 181–192. doi:10.1093/cercor/7.2.181

Essen, D. C., & Zeki, S. M. (1978). The topographic organization of rhesus monkey prestriate cortex. J Physiol, 277, 193–226. doi:10.1113/jphysiol 1978.sp012269

Feuerriegel, D., Churches, O., Hofmann, J., & Keage, H. A. D. (2015). The N170 and face perception in psychiatric and neurological disorders: A systematic review. Clin Neurophysiol, 126(6), 1141–1158. doi:10.1016/j.clinph.2014.09.015

Fischer, B., & Ramsperger, E. (1984). Human express saccades: extremely short reaction times of goal directed eye movements. Experimental Brain Research, 57(1), 191–195. doi:10.1007/BF00231145

Floris, D. L., Llera, A., Zabihi, M., Moessnang, C., Jones, E. J. H., Mason, L., Langer, N. (2025). A multimodal neural signature of face processing in autism within the fusiform gyrus. Nat Ment Health, 3(1), 31–45. doi:10.1038/s44220-024-00349-4

Flösch, K. P., Flaisch, T., Imhof, M. A., & Schupp, H. T. (2024). Alpha/beta oscillations reveal cognitive and affective brain states associated with role taking in a dyadic cooperative game. Cereb Cortex, 34(1). doi:10.1093/cercor/bhad487

Foster, J. J., & Awh, E. (2019). The role of alpha oscillations in spatial attention: limited evidence for a suppression account. Curr Opin Psychol, 29, 34–40. doi:10.1016/j.copsyc.2018.11.001

Foster, J. J., Sutterer, D. W., Serences, J. T., Vogel, E. K., & Awh, E. (2017). Alpha-Band Oscillations Enable Spatially and Temporally Resolved Tracking of Covert Spatial Attention. Psychol Sci, 28(7), 929–941. doi:10.1177/0956797617699167

Foxe, J. J., Simpson, G. V., & Ahlfors, S. P. (1998). Parieto-occipital ∼1 0Hz activity reflects anticipatory state of visual attention mechanisms. NeuroReport, 9(17), 3929–3933. Retrieved from https://journals.lww.com/neuroreport/fulltext/1998/12010/parieto_occipital 1_0h z_activity_reflects.30.aspx

Foxe, J. J., & Snyder, A. C. (2011). The Role of Alpha-Band Brain Oscillations as a Sensory Suppression Mechanism during Selective Attention. Front Psychol, 2, 154. doi:10.3389/fpsyg.2011.00154

Foxe, J. J., & Snyder, A. C. (2011). The Role of Alpha-Band Brain Oscillations as a Sensory Suppression Mechanism during Selective Attention. Frontiers in Psychology, 2. doi:10.3389/fpsyg.2011.00154

Garland, E. L., Froeliger, B., & Howard, M. O. (2015). Neurophysiological evidence for remediation of reward processing deficits in chronic pain and opioid misuse following treatment with Mindfulness-Oriented Recovery Enhancement: exploratory ERP findings from a pilot RCT. J Behav Med, 38(2), 327–336. doi:10.1007/s10865-014-9607-0

Greimel, E., Feldmann, L., Piechaczek, C., Oort, F., Bartling, J., Schulte-Ruther, M., & Schulte-Korne, G. (2020). Study protocol for a randomised-controlled study on emotion regulation training for adolescents with major depression: the KONNI study. BMJ Open, 10(9), e036093. doi:10.1136/bmjopen-2019-036093

Gross, J. J. (2007). *Handbook of Emotion Regulation*: Guilford Press.

Haegens, S., Nácher, V., Luna, R., Romo, R., & Jensen, O. (2011). α-Oscillations in the monkey sensorimotor network influence discrimination performance by rhythmical inhibition of neuronal spiking. Proceedings of the National Academy of Sciences, 108(48), 19377–19382. doi:doi:10.1073/pnas.1117190108

Hajcak, G., & Foti, D. (2020). Significance?& Significance! Empirical, methodological, and theoretical connections between the late positive potential and P300 as neural responses to stimulus significance: An integrative review. Psychophysiology, 57(7), e13570. doi:10.1111/psyp.13570

Halgren, M., Ulbert, I., Bastuji, H., Fabó, D., Erőss, L., Rey, M., . . . Cash, S. S. (2019). The generation and propagation of the human alpha rhythm. Proceedings of the National Academy of Sciences, 116(47), 23772–23782. doi:doi:10.1073/pnas.1913092116

Hertig-Godeschalk, A., Skorucak, J., Malafeev, A., Achermann, P., Mathis, J., & Schreier, D. R. (2019). Microsleep episodes in the borderland between wakefulness and sleep. Sleep, 43(1). doi:10.1093/sleep/zsz163

Huang, W. A., Stitt, I. M., Negahbani, E., Passey, D. J., Ahn, S., Davey, M., . . . Fröhlich, F. (2021). Transcranial alternating current stimulation entrains alpha oscillations by preferential phase synchronization of fast-spiking cortical neurons to stimulation waveform. Nature Communications, 12(1), 3151. doi:10.1038/s41467-021-23021-2

Hubel, D. H., & Wiesel, T. N. (2004). Brain and Visual Perception: The Story of a 25-year Collaboration: Oxford University Press.

Iwasaki, M., Kellinghaus, C., Alexopoulos, A. V., Burgess, R. C., Kumar, A. N., Han, Y. H., . . . Leigh, R. J. (2005). Effects of eyelid closure, blinks, and eye movements on the electroencephalogram. Clinical Neurophysiology, 116(4), 878–885. 10.1016/j.clinph.2004.11.001

Jensen, O. (2024). Distractor inhibition by alpha oscillations is controlled by an indirect mechanism governed by goal-relevant information. Communications Psychology, 2(1), 36. doi:10.1038/s44271-024-00081-w

Jensen, O., & Mazaheri, A. (2010). Shaping functional architecture by oscillatory alpha activity: gating by inhibition. Front Hum Neurosci, 4, 186. doi:10.3389/fnhum.2010.00186

Johnston, P., Molyneux, R., & Young, A. W. (2014). The N170 observed ‘in the wild’: robust event-related potentials to faces in cluttered dynamic visual scenes. Social Cognitive and Affective Neuroscience, 10(7), 938–944. doi:10.1093/scan/nsu136

Jones, S. A., Cowper-Smith, C. D., & Westwood, D. A. (2014). Directional interactions between current and prior saccades. Front Hum Neurosci, 8, 872. doi:10.3389/fnhum.2014.00872

Kala, S., Rolison, M. J., Trevisan, D. A., Naples, A. J., Pelphrey, K., Ventola, P., & McPartland, J. C. (2021). Brief Report: Preliminary Evidence of the N170 as a Biomarker of Response to Treatment in Autism Spectrum Disorder. Front Psychiatry, 12, 709382. doi:10.3389/fpsyt.2021.709382

Kastner, S., Pinsk, M. A., De Weerd, P., Desimone, R., & Ungerleider, L. G. (1999). Increased Activity in Human Visual Cortex during Directed Attention in the Absence of Visual Stimulation. Neuron, 22(4), 751–761. 10.1016/S0896-6273(00)80734-5

Keating, E. G., & Gooley, S. G. (1988). Saccadic disorders caused by cooling the superior colliculus or the frontal eye field, or from combined lesions of both structures. Brain Res, 438(1-2), 247–255. doi:10.1016/0006-8993(88)91343-1

Keren, A. S., Yuval-Greenberg, S., & Deouell, L. Y. (2010). Saccadic spike potentials in gamma-band EEG: characterization, detection and suppression. Neuroimage, 49(3), 2248–2263. doi:10.1016/j.neuroimage.2009.10.057

Kingstone, A., & Klein, R. M. (1993). What are human express saccades? Percept Psychophys, 54(2), 260–273. doi:10.3758/bf03211762

Kirchner, J., Watson, T., Bauer, J., & Lappe, M. (2022). All the ways your eyes can move if you don’t look: Eyeball lifting, retraction and compression during blinks. bioRxiv, 2022.2005.2011.491482. doi:10.1101/2022.05.11.491482

Klimesch, W. (2012). Alpha-band oscillations, attention, and controlled access to stored information. Trends in Cognitive Sciences, 16(12), 606–617. 10.1016/j.tics.2012.10.007

Kornrumpf, B., Dimigen, O., & Sommer, W. (2017). Lateralization of posterior alpha EEG reflects the distribution of spatial attention during saccadic reading. Psychophysiology, 54(6), 809–823. 10.1111/psyp.12849

Kurki, I., Hyvärinen, A., & Henriksson, L. (2022). Dynamics of retinotopic spatial attention revealed by multifocal MEG. Neuroimage, 263, 119643. 10.1016/j.neuroimage.2022.119643

Lakens, D. (2022). Sample Size Justification. Collabra: Psychology, 8(1). doi:10.1525/collabra.33267

Lang, P. J. (1979). A Bio-Informational Theory of Emotional Imagery. Psychophysiology, 16(6), 495–512. 10.1111/j.1469-8986.1979.tb01511.x

Lerner, Y., Hendler, T., Malach, R., Harel, M., Leiba, H., Stolovitch, C., & Pianka, P. (2006). Selective fovea-related deprived activation in retinotopic and high-order visual cortex of human amblyopes. Neuroimage, 33(1), 169–179. doi:10.1016/j.neuroimage.2006.06.026

Liu, B., Nobre, A. C., & van Ede, F. (2022). Functional but not obligatory link between microsaccades and neural modulation by covert spatial attention. Nat Commun, 13(1), 3503. doi:10.1038/s41467-022-31217-3

Liu, B., Nobre, A. C., & van Ede, F. (2023). Microsaccades transiently lateralise EEG alpha activity. Prog Neurobiol, 224, 102433. doi:10.1016/j.pneurobio.2023.102433

Luck, S. J. (1995). Multiple mechanisms of visual-spatial attention: recent evidence from human electrophysiology. Behavioural Brain Research, 71(1), 113–123. 10.1016/0166-4328(95)00041-0

Majaj, N. J., Carandini, M., & Movshon, J. A. (2007). Motion integration by neurons in macaque MT is local, not global. J Neurosci, 27(2), 366–370. doi:10.1523/JNEUROSCI.3183-06.2007

Marder, M. A., & Miller, G. A. (2024). The future of psychophysiology, then and now. Biol Psychol, 189, 108792. doi:10.1016/j.biopsycho.2024.108792

Maris, E., & Oostenveld, R. (2007). Nonparametric statistical testing of EEG- and MEG-data. Journal of Neuroscience Methods, 164(1), 177–190. 10.1016/j.jneumeth.2007.03.024

Martinez-Conde, S., Krauzlis, R., Miller, J., Morrone, C., Williams, D., & Kowler, E. (2008). Eye movements and the perception of a clear and stable visual world. J Vis, 8(14), 1. doi:10.1167/8.14.i

Martinez-Conde, S., Macknik, S. L., & Hubel, D. H. (2004). The role of fixational eye movements in visual perception. Nat Rev Neurosci, 5(3), 229–240. doi:10.1038/nrn1348

Mason, L., Moessnang, C., Chatham, C., Ham, L., Tillmann, J., Dumas, G., Jones, E. J. H. (2022). Stratifying the autistic phenotype using electrophysiological indices of social perception. Science Translational Medicine, 14(658), eabf8987. doi:doi:10.1126/scitranslmed.abf8987

Maunsell, J. H., & Van Essen, D. C. (1987). Topographic organization of the middle temporal visual area in the macaque monkey: representational biases and the relationship to callosal connections and myeloarchitectonic boundaries. J Comp Neurol, 266(4), 535–555. doi:10.1002/cne.902660407

Meyberg, S., Werkle-Bergner, M., Sommer, W., & Dimigen, O. (2015). Microsaccade-related brain potentials signal the focus of visuospatial attention. Neuroimage, 104, 79–88. 10.1016/j.neuroimage.2014.09.065

Miller, G. A. (2010). Mistreating Psychology in the Decades of the Brain. Perspectives on Psychological Science, 5(6), 716–743. doi:10.1177/1745691610388774

Movshon, J. A., & Newsome, W. T. (1996). Visual response properties of striate cortical neurons projecting to area MT in macaque monkeys. J Neurosci, 16(23), 7733–7741. doi:10.1523/JNEUROSCI.16-23-07733.1996

Oostenveld, R., Fries, P., Maris, E., & Schoffelen, J.-M. (2011). FieldTrip: Open Source Software for Advanced Analysis of MEG, EEG, and Invasive Electrophysiological Data. Computational Intelligence and Neuroscience, 2011(1), 156869. 10.1155/2011/156869

Peirce, J., Gray, J. R., Simpson, S., MacAskill, M., Höchenberger, R., Sogo, H., . . . Lindeløv, J. K. (2019). PsychoPy2: Experiments in behavior made easy. Behavior Research Methods, 51(1), 195–203. doi:10.3758/s13428-018-01193-y

Popov, T., Gips, B., Weisz, N., & Jensen, O. (2023). Brain areas associated with visual spatial attention display topographic organization during auditory spatial attention. Cereb Cortex, 33(7), 3478–3489. doi:10.1093/cercor/bhac285

Popov, T., Miller, G. A., Rockstroh, B., Jensen, O., & Langer, N. (2021). Alpha oscillations link action to cognition: An oculomotor account of the brain’s dominant rhythm. bioRxiv, 2021.2009.2024.461634. doi:10.1101/2021.09.24.461634

Popov, T., Miller, G. A., Rockstroh, B., & Weisz, N. (2013). Modulation of α power and functional connectivity during facial affect recognition. J Neurosci, 33(14), 6018–6026. doi:10.1523/jneurosci.2763-12.2013

Popov, T., & Staudigl, T. (2023). Cortico-ocular coupling in the service of episodic memory formation. Prog Neurobiol, 227, 102476. doi:10.1016/j.pneurobio.2023.102476

Popov, T., Steffen, A., Weisz, N., Miller, G. A., & Rockstroh, B. (2012). Cross-frequency dynamics of neuromagnetic oscillatory activity: two mechanisms of emotion regulation. Psychophysiology, 49(12), 1545–1557. doi:10.1111/j.1469-8986.2012.01484.x

Popov, T., Steffen, A., Weisz, N., Miller, G. A., & Rockstroh, B. (2012). Cross-frequency dynamics of neuromagnetic oscillatory activity: Two mechanisms of emotion regulation. Psychophysiology, 49(12), 1545–1557. 10.1111/j.1469-8986.2012.01484.x

Popov, T., & Szyszka, P. (2020). Alpha oscillations govern interhemispheric spike timing coordination in the honey bee brain. Proc Biol Sci, 287(1921), 20200115. doi:10.1098/rspb.2020.0115

Popov, T., Trondle, M., Baranczuk-Turska, Z., Pfeiffer, C., Haufe, S., & Langer, N. (2023). Test-retest reliability of resting-state EEG in young and older adults. Psychophysiology, 60(7), e14268. doi:10.1111/psyp.14268

Popov, T. G., Rockstroh, B. S., Popova, P., Carolus, A. M., & Miller, G. A. (2014). Dynamics of alpha oscillations elucidate facial affect recognition in schizophrenia. Cogn Affect Behav Neurosci, 14(1), 364–377. doi:10.3758/s13415-013-0194-2

Popova, P., Popov, T. G., Wienbruch, C., Carolus, A. M., Miller, G. A., & Rockstroh, B. S. (2014). Changing facial affect recognition in schizophrenia: effects of training on brain dynamics. Neuroimage Clin, 6, 156–165. doi:10.1016/j.nicl.2014.08.026

Powers, W. T. (1973). Behavior: The control of perception. Oxford, England: Aldine.

Quax, S. C., Dijkstra, N., van Staveren, M. J., Bosch, S. E., & van Gerven, M. A. J. (2019). Eye movements explain decodability during perception and cued attention in MEG. Neuroimage, 195, 444–453. doi:10.1016/j.neuroimage.2019.03.069

Redding, Z. V., & Fiebelkorn, I. C. (2023). Distractor suppression does and does not depend on pre-distractor alpha-band activity. bioRxiv. doi:10.1101/2023.07.18.549512

Robinson, D. A. (1972). Eye movements evoked by collicular stimulation in the alert monkey. Vision Research, 12(11), 1795–1808. 10.1016/0042-6989(72)90070-3

Saalmann, Y. B., Pinsk, M. A., Wang, L., Li, X., & Kastner, S. (2012). The pulvinar regulates information transmission between cortical areas based on attention demands. Science, 337(6095), 753–756. doi:10.1126/science.1223082

Schiller, P. H., Haushofer, J., & Kendall, G. (2004). An examination of the variables that affect express saccade generation. Vis Neurosci, 21(2), 119–127. doi:10.1017/s0952523804042038

Schiller, P. H., Sandell, J. H., & Maunsell, J. H. (1987). The effect of frontal eye field and superior colliculus lesions on saccadic latencies in the rhesus monkey. J Neurophysiol, 57(4), 1033–1049. doi:10.1152/jn.1987.57.4.1033

Schiller, P. H., & Stryker, M. (1972). Single-unit recording and stimulation in superior colliculus of the alert rhesus monkey. J Neurophysiol, 35(6), 915–924. doi:10.1152/jn.1972.35.6.915

Schiller, P. H., Stryker, M., Cynader, M., & Berman, N. (1974). Response characteristics of single cells in the monkey superior colliculus following ablation or cooling of visual cortex. J Neurophysiol, 37(1), 181–194. doi:10.1152/jn.1974.37.1.181

Schiller, P. H., & Tehovnik, E. J. (2001). Chapter 9 Look and see: how the brain moves your eyes about. In Progress in Brain Research (Vol. 134, pp. 127–142): Elsevier.

Schiller, P. H., & Tehovnik, E. J. (2001). Look and see: how the brain moves your eyes about. Prog Brain Res, 134, 127–142. doi:10.1016/s0079-6123(01)34010-4

Schiller, P. H., & Tehovnik, E. J. (2003). Cortical inhibitory circuits in eye-movement generation. Eur J Neurosci, 18(11), 3127–3133. doi:10.1111/j.1460-9568.2003.03036.x

Schiller, P. H., & Tehovnik, E. J. (2005). Neural mechanisms underlying target selection with saccadic eye movements. Prog Brain Res, 149, 157–171. doi:10.1016/s0079-6123(05)49012-3

Schubring, D., Kraus, M., Stolz, C., Weiler, N., Keim, D. A., & Schupp, H. (2020). Virtual Reality Potentiates Emotion and Task Effects of Alpha/Beta Brain Oscillations. Brain Sciences, 10(8), 537. Retrieved from https://www.mdpi.com/2076-3425/10/8/537

Schubring, D., & Schupp, H. T. (2019). Affective picture processing: Alpha- and lower beta-band desynchronization reflects emotional arousal. Psychophysiology, 56(8), e13386. 10.1111/psyp.13386

Schubring, D., & Schupp, H. T. (2020). Emotion and Brain Oscillations: High Arousal is Associated with Decreases in Alpha- and Lower Beta-Band Power. Cerebral Cortex, 31(3), 1597–1608. doi:10.1093/cercor/bhaa312

Schupp, H. T., Cuthbert, B. N., Bradley, M. M., Cacioppo, J. T., Ito, T., & Lang, P. J. (2000). Affective picture processing: the late positive potential is modulated by motivational relevance. Psychophysiology, 37(2), 257–261.

Seiple, W., Rosen, R. B., & Garcia, P. M. T. (2013). Abnormal Fixation in Individuals With Age-Related Macular Degeneration When Viewing an Image of a Face. Optometry and Vision Science, 90(1). Retrieved from https://journals.lww.com/optvissci/fulltext/2013/01000/abnormal_fixation_in_individuals_with_age_related.10.aspx

Skorucak, J., Hertig-Godeschalk, A., Schreier, D. R., Malafeev, A., Mathis, J., & Achermann, P. (2019). Automatic detection of microsleep episodes with feature-based machine learning. Sleep, 43(1). doi:10.1093/sleep/zsz225

Spiering, L., & Dimigen, O. (2024). (Micro)saccade-related potentials during face recognition: A study combining EEG, eye-tracking, and deconvolution modeling. *Attention, Perception*, & Psychophysics. doi:10.3758/s13414-024-02846-1

Stine, G. M., Trautmann, E. M., Jeurissen, D., & Shadlen, M. N. (2023). A neural mechanism for terminating decisions. Neuron, 111(16), 2601–2613.e2605. 10.1016/j.neuron.2023.05.028

Tehovnik, E. J., Slocum, W. M., Carvey, C. E., & Schiller, P. H. (2005). Phosphene induction and the generation of saccadic eye movements by striate cortex. J Neurophysiol, 93(1), 1–19. doi:10.1152/jn.00736.2004

Tullis, T., & Albert, B. (2013). Chapter 7 - Behavioral and Physiological Metrics. In T. Tullis & B. Albert (Eds.), Measuring the User Experience (Second Edition) (pp. 163–186). Boston: Morgan Kaufmann.

Van Essen, D. C., Newsome, W. T., & Maunsell, J. H. (1984). The visual field representation in striate cortex of the macaque monkey: asymmetries, anisotropies, and individual variability. Vision Res, 24(5), 429–448. doi:10.1016/0042-6989(84)90041-5

Vogel, E. K., & Luck, S. J. (2000). The visual N1 component as an index of a discrimination process. Psychophysiology, 37(2), 190–203.

Wang, L., Mruczek, R. E., Arcaro, M. J., & Kastner, S. (2015). Probabilistic Maps of Visual Topography in Human Cortex. Cereb Cortex, 25(10), 3911–3931. doi:10.1093/cercor/bhu277

Ward, A. S. J. M. (2017, 06 Dec 2017). Here’s What You Can Learn About Architecture from Tracking People’s Eye Movements. ArchDaily. Retrieved from https://www.archdaily.com/884945/heres-what-you-can-learn-about-architecture-from-tracking-peoples-eye-movements

Westner, B. U., & Dalal, S. S. (2019). Faster than the brain’s speed of light: Retinocortical interactions differ in high frequency activity when processing darks and lights. bioRxiv, 153551. doi:10.1101/153551

Westner, B. U., Lubell, J. I., Jensen, M., Hokland, S., & Dalal, S. S. (2021). Contactless measurements of retinal activity using optically pumped magnetometers. Neuroimage, 243, 118528. doi:10.1016/j.neuroimage.2021.118528

Woldorff, M. G., Fox, P. T., Matzke, M., Lancaster, J. L., Veeraswamy, S., Zamarripa, F., . . . Jerabek, P. (1997). Retinotopic organization of early visual spatial attention effects as revealed by PET and ERPs. Human Brain Mapping, 5(4), 280–286. 10.1002/(SICI)1097-0193(1997)5:4<280::AID-HBM13>3.0.CO;2-I

Worden, M. S., Foxe, J. J., Wang, N., & Simpson, G. V. (2000). Anticipatory biasing of visuospatial attention indexed by retinotopically specific alpha-band electroencephalography increases over occipital cortex. J Neurosci, 20(6), Rc63. doi:10.1523/JNEUROSCI.20-06-j0002.2000

Wright, K. W. (2006). Anatomy and Physiology of Eye Movements. In K. W. Wright, P. H. Spiegel, & L. S. Thompson (Eds.), Handbook of Pediatric Strabismus and Amblyopia (pp. 24–69). New York, NY: Springer New York.

Yarbus, A. L. (2013). Eye movements and vision: Springer.

Yuval-Greenberg, S., Tomer, O., Keren, A. S., Nelken, I., & Deouell, L. Y. (2008). Transient induced gamma-band response in EEG as a manifestation of miniature saccades. Neuron, 58(3), 429–441. doi:10.1016/j.neuron.2008.03.027

Zhu, J., Zhou, X. M., Constantinidis, C., Salinas, E., & Stanford, T. R. (2024). Parallel signatures of cognitive maturation in primate antisaccade performance and prefrontal activity. iScience, 27(8), 110488. 10.1016/j.isci.2024.110488

Zsigo, C., Feldmann, L., Oort, F., Piechaczek, C., Bartling, J., Schulte-Rüther, M., . . . Greimel, E. (2024). Emotion regulation training for adolescents with major depression: Results from a randomized controlled trial. Emotion, 24(4), 975–991. doi:10.1037/emo0001328

